# Paralleled Dynamics of Arabidopsis Root Exudation and SynCom Assembly in a Controlled Environment

**DOI:** 10.64898/2026.01.29.702624

**Authors:** Charlotte Joller, Jan Waelchli, Joelle Schlaepfer, Klaus Schlaeppi

## Abstract

Plant roots host defined microbial communities that differ from those found in the surrounding soil and these communities shift dynamically in response to plant development and environmental changes. Whilst it is widely accepted that root exudates play a key role in the assembly and dynamics of root-associated microbial communities, the underlying mechanisms are not well understood. This is partly due to a lack of controlled experimental systems that monitor both exudate- and microbiome-dynamics simultaneously. Here, we compared two microcosm systems commonly used in either root microbiome (clay particle-based) or root exudate studies (glass bead-based) for their suitability to simultaneously monitor both aspects. We evaluated these systems based on plant performance, bacterial growth, and time-resolved community and exudate profiling. In both systems, we reveal an exudate effect, characterised by higher bacterial diversity and *Pseudomonas* abundances in proximity to plant roots. While clay particles impeded exudate recovery, even when plants were removed from microcosms for exudate collection, the glass bead set-up allowed us to uncover dynamic exudate shifts during bacterial community establishment. This highlighted a transient increase of glucosinolates upon root colonisation by initially dominant *Pseudomonas* species. Overall, the comparison proved only the glass bead-based semi-hydroponic system to be suitable for the paralleled study of exudate and root microbiome dynamics.

## Introduction

In natural environments, plant roots are colonised by a plethora of microorganisms that confer numerous benefits, enhancing plant resilience to both biotic and abiotic stresses (Beeckman and Eshel, 2024; Seenivasagan and Babalola, 2021). Root microbiomes are composed of core microbial taxa that consistently associate with plant roots across different climates, soil types and plant genotypes (Almario et al., 2022; Durán et al., 2022; Lundberg et al., 2021; Thiergart et al., 2020). Although it is well established that plants interact with the resident soil microbiota through the secretion of exudates, the precise mechanisms by which this contributes to shaping these defined root-associated microbial communities are largely elusive (Rasmann et al., 2024).

Deciphering the chemical dialogue between plant roots and associated microbiota presents a major challenge. Soil not only harbours a “daunting complexity of microbial communities” (Tecon et al., 2019) that metabolises exudates, but also presents a chemically diverse matrix and contains minerals that readily absorb metabolites, thereby interfering with the detection of plant-derived compounds (Aleklett et al., 2018; Oburger and Jones, 2018; Tecon et al., 2019). In addition, both root exudation and microbiomes undergo continuous and dynamic shifts. Beyond responses to biotic and abiotic stress, plant-associated microbiomes vary depending on the season and the developmental stage or time since inoculation (Almario et al., 2022; Bartoli et al., 2018; Beilsmith et al., 2021; Chaparro et al., 2014; Chen et al., 2024; Dibner et al., 2021; Mayer et al., 2025). Similarly, root exudation also changes with the plant developmental stage, diurnal time point or abiotic stress (Chaparro et al., 2013; McLaughlin et al., 2023b; Robert et al., 2025; Zhalnina et al., 2018). Recently, we have reported dynamic responses in exudates to the presence of microbe- and damage-associated molecular patterns (MAMPs, DAMPs) with a significant shift in exudation already measurable after four hours (Joller et al., 2024). Collectively, the physicochemical properties of the growth matrix, the presence and complexity of microbiota, and the requirement of repeated samplings are crucial challenges for monitoring the interactions between root exudates and microbiomes. To overcome these challenges, high-resolution and time-resolved experimental systems are needed (Kuijken et al., 2015; McLaughlin et al., 2025; Oburger and Jones, 2018; Salomonsen et al., 2024).

In the last decade, reductionist approaches have provided major advancements in the field of plant-microbe interactions. Generally, such approaches aim at deconstructing the complexity seen in natural environments to later reconstruct elements one by one under controlled laboratory conditions. The reductionist strategy makes it possible to establish causal links and has been instrumental in gaining mechanistic insights into plant-microbe interactions (Chesneau et al., 2025; Tecon et al., 2019). In the context of microbiome work, a first element of reductionist approaches is the use of a defined microbiota. For this synthetic communities (SynComs) have been developed (Durán et al., 2025; Jing et al., 2024; Li et al., 2024; Niu et al., 2017; Northen et al., 2024; Paredes et al., 2018; Vorholt et al., 2017). SynComs serve as reconstructed representative microbial consortia for natural communities that were previously deconstructed into culture collections (Bai et al., 2015; Gómez-Repollés et al., 2025; Thoenen et al., 2024, 2023; Wippel et al., 2021).

A second element of reductionist approaches are fully defined experimental systems. Such systems with a defined microbiota, referred to as gnotobiotic systems, are frequently used in combination with SynComs to study plant-microbe interactions. They range from very simple hydroponics to agar-based systems (Chen et al., 2024; Ma et al., 2022; Vogel et al., 2012; Wippel et al., 2021) to more complex systems mimicking soil conditions using chemically defined matrices such as clay, sand, vermiculate or artificial soils (Aleklett et al., 2018; Bai et al., 2015; Ghosheh et al., 2025; Kremer et al., 2021; Ma et al., 2022; Salomonsen et al., 2024). Finally, there are also very complex, custom-made, fully tractable and tuneable synthetic ecosystems (Schmidt et al., 2021). Many of these systems are used to study microbial contributions to plant growth and defence (Castrillo et al., 2017; Finkel et al., 2020; Vogel et al., 2012).

However, root exudates and microbiomes are rarely monitored in the same system, which is essential to establish causal relationships between exudation and microbiome dynamics. The main reason is that, whilst most microbiome researchers aim at maintaining a soil-like environment, exudate metabolites are hard to trace in complex chemical backgrounds (Ma et al., 2022; Salomonsen et al., 2024; Sasse et al., 2020). Therefore, exudates are usually sampled from plants cultured hydroponically, or in hybrid approaches where plants are first cultured in soil and then transferred to hydroponics for exudate collection (Oburger and Jones, 2018). Notable examples of studies where exudate and microbiome composition have been measured in parallel, thus also mostly relied on hydroponic cultivation (Korenblum et al., 2020; Salomonsen et al., 2024).

Systematic comparisons of reductionist systems, commonly used to study either root exudates or microbiomes, are lacking when it comes to the paralleled analysis of both root exudate and microbiome dynamics. Here, we compared two such systems using the same SynCom for plant performance, time-resolved root exudate profiles and recovery, along with bacterial performance and community composition: a semi-hydroponic system incorporating glass beads as a physical support (glass-microcosms, McLaughlin et al., 2023a), rather than a fully liquid hydroponic system, was compared with a soil-like system in which clay particles provided physical support (clay-microcosms, Bai et al., 2015). Experiments were conducted using *Arabidopsis thaliana* (hereafter Arabidopsis), which, is an established model organism in plant-microbiome research (Poupin et al., 2023). We show that while both systems support plant growth well and present comparable root bacterial community densities and structures, the clay matrix absorbs a large proportion of the exudate signal, whereas the glass-microcosms enable time-resolved recovery of diverse metabolites. The semi-hydroponic glass-microcosms proved to be the suitable system for simultaneous exudate – microbiome analysis. As proof-of-concept, we show the time-resolved and highly reproducible assembly of root bacterial communities and the associated changes in exudate chemistry. Most notably, we found an increasing diverse consortia alongside a transient increase of aliphatic glucosinolate levels.

## Results

### Similar plant performance and bacterial community assembly in glass- and clay-microcosms

For simultaneous analysis of exudates and microbiomes, we evaluated glass- and clay-microcosms with gnotobiotic Arabidopsis (**Fig. 1A**). Plants were transferred as sterile seedlings to microcosms and inoculated with a synthetic community composed of nine strains (SynCom9, **Table S1**). These strains were originally isolated from roots of Arabidopsis cultured in clay microcosms with a soil extract and were chosen to represent the most abundant strains while also preserving some of the phylogenetic diversity (Joller et al., in preperation).

**Fig. 1.**
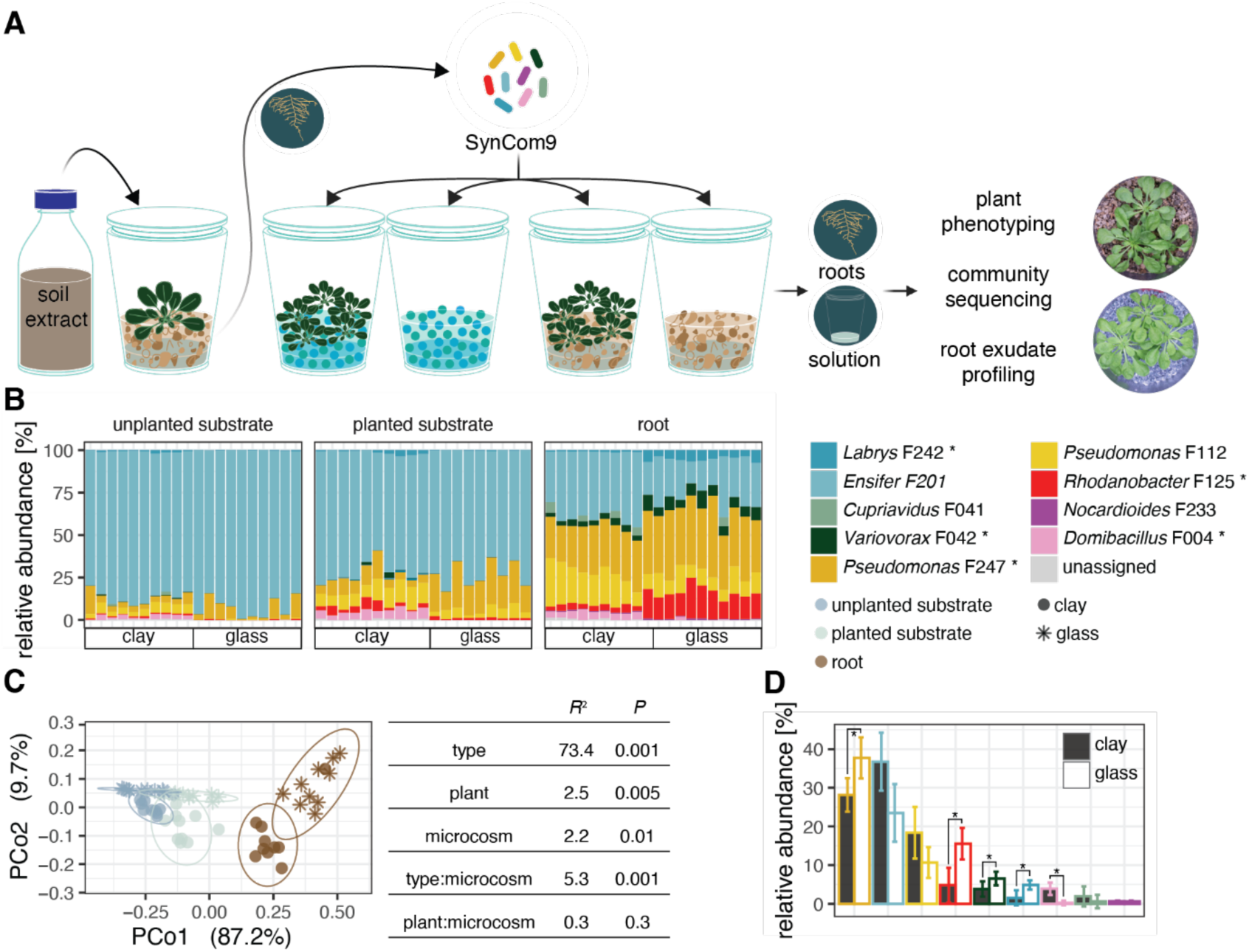
SynCom9 communities assemble similarly in substrate and on roots in glass- and clay-microcosms. **A** Graphical overview of the experimental setup in gnotobiotic glass- (blue beads) and clay (brownish flakes) microcosms used to evaluate their suitability for paralleled microbiome and exudate profiling. Three to five plants were cultured axenically or with a mini SynCom comprising 9 bacterial strains (SynCom9) isolated from Arabidopsis. Unplanted and planted conditions were compared to determine the influence of plants on metabolite and microbiome profiles in root-surrounding solutions. **B-C** Root and substrate SynCom9 communities in glass-and clay-microcosms after 2 weeks of co-cultivation (N = 9-10 microcosms à 3 plants each). **B** Relative abundances of SynCom9 strains. **C** Bray-Curtis dissimilarities of communities displayed in a Principal Coordinate Analysis (PCoA). The table shows PERMANOVA results (type: substrate vs root, plant: unplanted vs planted, microcosm: glass vs clay). Ellipses represent 95% confidence intervals. **D** Differential abundance analysis of SynCom9 members in roots. The graph shows mean relative abundances and standard deviations of each strain. Differential abundance between glass and clay microcosms was determined via aldex2 (Fernandes et al., 2012), acombc (Lin & Peddada, 2020), maaslin2 (Mallick et al., 2021) and metagenomeSeq (Paulson et al., 2013). Strains that were designated differentially abundant by at least three of the methods are marked with an asterisk.

Plants grew well in both glass and clay matrices as evident from similar root and shoot biomass after two weeks in microcosms and they did not exhibit negative phenotypes like developmental anomalies or disease development (**Fig. S1**). Root biomass was 30-45% lower in glass-compared to clay-microcosms but there was no difference in shoot biomass. There was no effect of inoculation with the SynCom9 on root or shoot weights in either microcosm type.

We assessed SynCom9 growth in the substrate solution and on roots based on colony-forming units per millilitre substrate solution and gram fresh weight, respectively. On roots, there was no difference in bacterial densities between glass- and clay microcosms (∼ 1 × 10^5^ CFU mg^-1^, **Fig. S2A**) and bacterial growth dynamics in substrate solutions followed the same trajectories in both microcosm types, with bacterial numbers plateauing 4 days after inoculation (dai) at ∼1–8 × 10^7^ CFU mL^-1^ (**Fig. S2B**). Of note, substrate samples were compared between planted and unplanted control microcosms to test for an *exudate effect* (i.e., higher bacterial numbers in substrate solutions of planted compared to unplanted microcosms). Only a small *exudate effect* and only significant in glass-microcosms was found (area under the curve (AUC) of CFU mL^-1^ over time; **Fig. S2C**).

To further understand bacterial community establishment in the two growth matrices, we measured SynCom9 composition using 16S rRNA gene amplicon sequencing. Alpha-diversity in substrate solutions was lower in glass-compared to clay-microcosm, while on roots, there was no difference in alpha-diversity between the two microcosms (**Fig. S2D**). In both microcosms, we observed the SynCom9 to increase in diversity with increasing influence of the root (from unplanted and planted substrates to roots). This was consistent with the sample-type specific community compositions (**Fig. 1B**) and the variation partitioning in ordination analysis (PCo1 largely separated the sample types and explained 73.4% of variation, **Fig. 1C**).

At the level of individual SynCom strains, the substrate communities were dominated by *Ensifer* F201 (∼90% and ∼70% of relative abundances in unplanted and planted microcosms, respectively), while the most abundant strains on roots were the two *Pseudomonas* strains (F112, F247). Next to the clear differences between sample types, SynCom9 community composition also differed between microcosms (7.5% of variation). In root communities, differential abundance analysis revealed significant effects for 5 out of the 9 SynCom9 strains: *Pseudomonas* F247, *Rhodanobacter* F125, *Variovorax* F042 and *Labrys* F242 being more abundant in glass-, and *Domibacillus* F004 more abundant in clay-microcosms (**Fig. 1D**). Note that *Domibacillus* F004 could barely be detected in glass-microcosms but was present throughout all clay-microcosm samples at low relative abundances.

In summary, the two microcosm types showed similar patterns in plant performance, bacterial growth, and community composition. In particular, each showed higher bacterial diversity and relative abundances of *Pseudomonas* in root-associated compared to substrate communities.

### Exudate signals absorbed in clay-microcosms

In the next step, we compared the two microcosms for their suitability to profile root exudates. Two approaches were employed for time-resolved exudate collection (2h intervals) from axenic plants grown in either glass- or clay-microcosms: first, *in situ*, with the plants left in microcosms, and subsequently, *ex situ*, with the plants removed from microcosms for exudate collection with a low metabolic background (**Fig. 2A**). To reduce the metabolic background of clay-microcosms “washed clay”-microcosms were included in the experiment (see *Materials and Methods*). Root exudate samples were analysed by flow-injection-analysis mass spectrometry (FIA MS, 2135 metabolites detected in this run). After background subtraction, 715 exudate metabolites were retained for analysis.

**Fig. 2.**
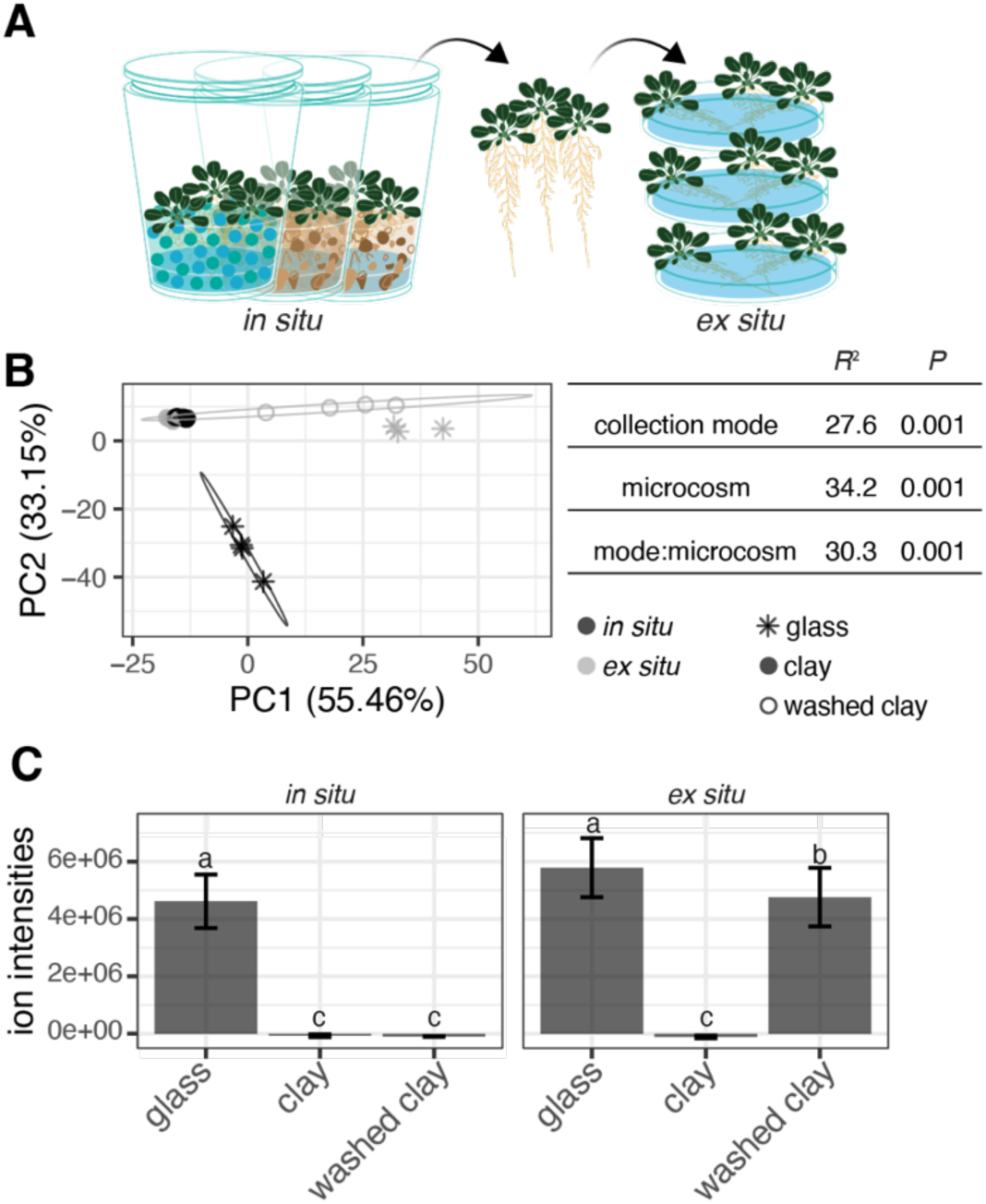
The clay matrix absorbs exudate signals even in *ex situ* exudate collection. Untargeted metabolomics analysis (flow-injection-analysis mass spectrometry (FIA MS) of (**A**) Arabidopsis exudates collected from glass- and clay-microcosms *in* and *ex situ*. **B** Principal Component Analysis (PCA) of normalised intensities of filtered metabolites (715 metabolites). Displayed are exudates collected *in situ* (black) and *ex situ* (grey) from glass- (asterisk symbols), clay- (circles) and washed clay-microcosms (open circles, N = 3-4 microcosms per condition comprising 5 plants each). The table shows PERMANOVA results calculated using Euclidean distances. Ellipses represent 95% confidence intervals. **C** Barplots showing mean summed metabolite intensities with the standard deviation. Displayed values are normalised by corresponding blank summed ion intensities (intensity_sample_ – mean intensity_blank_). Letters indicate significance as determined by ANOVA and Tukey’s HSD (N = 3-4).

Both, exudate collection mode and microcosm type had a significant effect on recovered and normalised exudate profiles (**Fig. 2B**). Exudate profiles of clay-grown plants overlapped with the blank profiles of the clay matrix (**Fig. S3A**). The only exception to this were *ex situ* collected exudates of plants grown in washed clay (**Fig. 2B, S3A**). Quantitatively, root exudates from glass-microcosms and *ex situ-*collected, washed clay-microcosms exhibited total summed ion intensities of ∼5 × 10⁶, whereas the other conditions resulted in near-zero ion intensities (**Fig. 2C**). These results show clearly that a large portion of root-derived compounds were absorbed by the clay matrix, even after *ex situ* collection.

Glass-microcosms showed a clear separation between ex situ and in situ exudate profiles. Indeed, 57% of all metabolites of various chemical classes significantly differed in their abundance between the two collection modes (37% more abundant and 20% less abundant in ex situ collected root exudates, Fig. S3B-D). Thus, differences between in situ and ex situ collected exudate profiles affected a substantial proportion of all detected metabolites and diverse metabolic classes.

Two main conclusions stem from this analysis: (i) sampling of root exudates from Arabidopsis plants grown in clay-microcosms was not possible, possibly hindered through metabolite absorption by the clay matrix. (ii) An *ex situ* collection led to marked differences in exudate profiles compared to *in situ* measurements, where plants are not removed from the matrix. Hence, only the *in situ* collection of exudates in the glass-microcosms is suitable for non-invasive and accurate exudate studies.

### Simultaneous analysis of metabolome and microbiome dynamics in glass- microcosms

Next, we used the glass-microcosms and tested the simultaneous analysis of exudate and microbiome dynamics over time. We followed SynCom9 root community establishment and the accompanying exudate changes in a time-course of early (0, 1, 2 and 4 days after inoculation, dai) to a late time point (12 dai,(**Fig. 3A**). The dense early timepoints were chosen as a first pilot revealed little change in SynCom9 composition after 4 dai (**Fig. S4**). Plant performance was scored by non-invasive imaging of rosette area and root and shoot biomass were obtained from staggered destructive samplings. Consistent with the earlier comparison of matrices (**Fig. S1**), inoculation with the SynCom9 did not impact plant shoot fresh weights (**Fig. S5**). The same was observed for rosette area. Only at the latest time point, root fresh weights of SynCom inoculated plants were significantly lower than those of sterile plants.

**Fig. 3.**
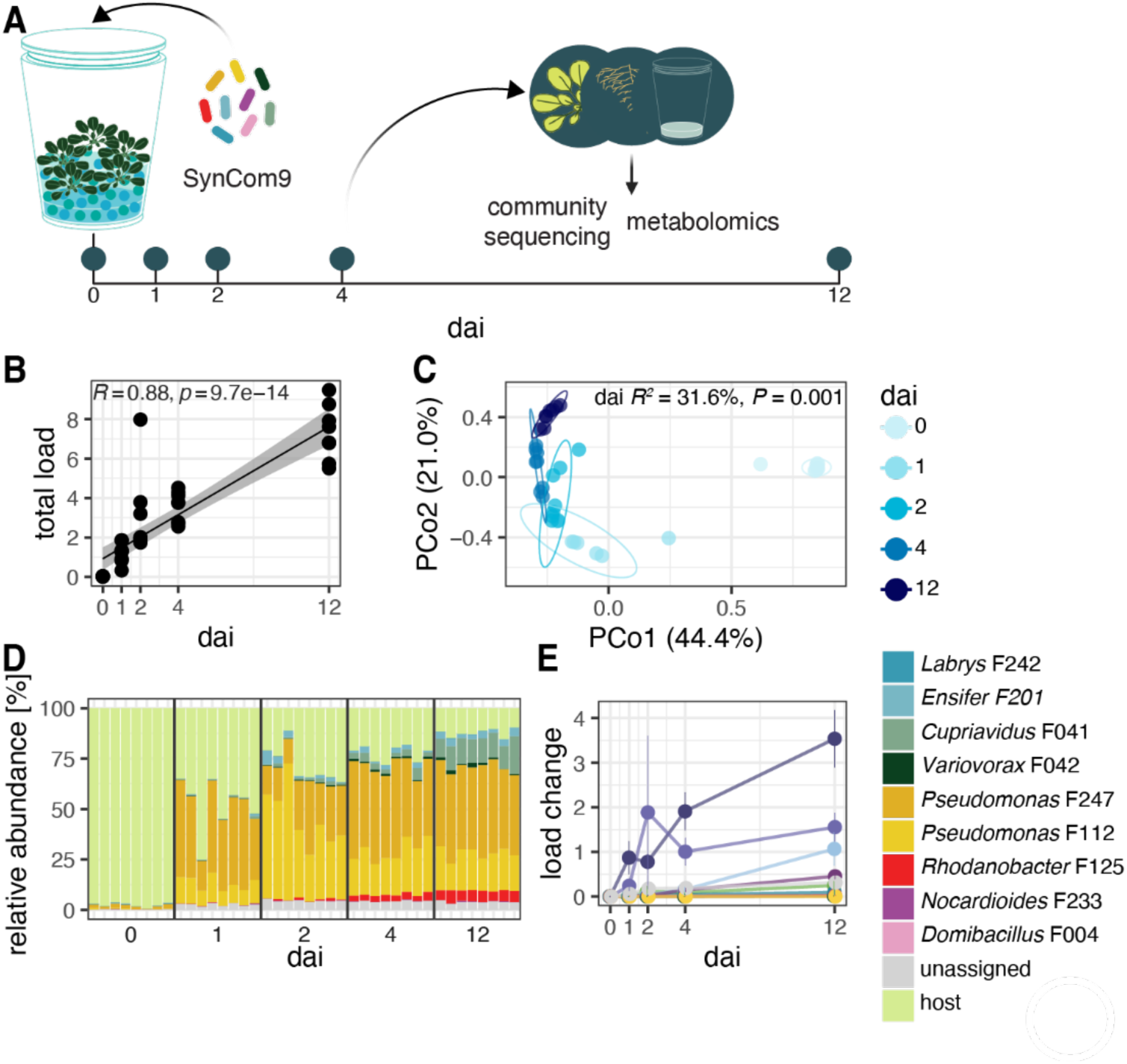
**Steady increase of bacterial load and diversity in SynCom9 root communities of plants cultured in glass- microcosms**. Root microbiome structure and bacterial load in glass-microcosms at 0, 1, 2, 4 and 12 days after inoculation (dai) determined by host-associated microbe PCR (hamPCR, Lundberg et al., 2021) (N = 8 microcosms per time point à 3 plants each). The bacterial load corresponds to the ratio of bacterial to host reads. **A** Experimental overview. **B** Pearson correlation of the total bacterial load with time. The grey area represents 95% confidence intervals. The correlation coefficient and *p-*value are indicated on the plot. **C** PCoA of Bray-Curtis dissimilarities between SynCom9 communities at different time points (light colours: early time points, dark colours: late time points) based on bacterial loads with PERMANOVA results. Ellipses represent 95% confidence intervals. **D** Relative abundances of SynCom9 strains and host reads in root samples. **E** Bacterial load of individual SynCom9 strains over time. The load for each time point is shown as the mean value with the standard deviation.

### Root communities became more even and diverse over time

We determined composition of SynCom9 on roots and their bacterial load via host-associated microbe PCR (hamPCR, Lundberg et al., 2021). The bacterial load increased steadily without reaching a plateau over the 12-day experimental period (**Fig. 3B**). This finding was confirmed by a strong correlation with root CFU counts (**Fig. S6A**). The composition of inoculated SynCom9 members matured over time, i.e. initially low abundant members became more abundant with time (indicated by Shannon diversity index, **Fig. S6B**). Secondly, the composition became less variable with time (Bray-Curtis dissimilarities **Fig. 3C, D**). As in the previous experiment (**Fig. S1**), the *Pseudomonas* strains dominated root communities, establishing early and reaching the highest densities (**Fig. 3D, E**). The *Pseudomonas* dynamics followed a conserved pattern between replicates. Whilst *Pseudomonas* F247 was dominant at time points 1, 4 and 12, *Pseudomonas* F112 was the most abundant strain at time point two. Except *Ensifer* F201, which showed an early peak similar to *Pseudomonas* F112, the other strains increased over time. *Domibacillus* F004 remained barely detectable across all timepoints, confirming its poor establishment in the glass matrix (**Fig. 3D, E, S7**). Together, these results point to a reproducible assembly of the SynCom9 on Arabidopsis roots in glass-microcosms, marked by early *Pseudomonas* dominance and a gradual convergence towards mature, less variable and more diverse communities with time.

### Inoculation with SynCom9 leads to increased exudation of glucosinolates

Alongside the microbiome profiling, we conducted untargeted metabolomics on exudates, roots, and shoots to characterise the plant metabolic responses to inoculation over time. Since in exudates of inoculated plants, plant-derived signals cannot be distinguished from bacteria-derived signals, we used tissue metabolic profiles to search for robust changes of plant metabolites. After background subtraction, 123, 975 and 838 metabolites were identified in exudate, roots and shoots, respectively.

Exudates roots and shoots differed markedly in their chemical composition (**Fig. S8**). Exudates were enriched in organic acids (41%), including amino acids, peptides, and analogues (AA, 26.4%) and tricarboxylic acids (TCA, 12.1%), compared to roots (12%, AA 8%, TCA 2%) and shoots (21%, AA 17%, TCA 1%). Roots and shoots contained more phenylpropanoids, especially flavonoids (21% and 14% vs. 2.5% in exudates). Organic oxygen compounds (e.g. alcohols, carbohydrates, carbohydrate conjugates or carbonyl compounds) were consistent across tissues (20–25%), but glucosinolates were more abundant in roots and shoots (14% and 16%) than in exudates (3.5%), which instead had higher levels of other organooxygen compounds (15.2%) and carbohydrates (7.2%). Exudate composition largely recapitulated that from the previous experiment, demonstrating the reproducibility of recovered exudate chemistry (**Fig. S8**). Time was the main driver of metabolic variation in all sample types (11% exudates, 20% roots, 17% shoots; **Fig. S9–S11**). For instance, glucosinolates increased and flavonoids decreased over time in roots and shoots (**Fig. 4A, S12**), while in exudates tricarboxylic acids were subjected to a stronger temporal shift (decreasing from 17% to 9%, **Fig. 4A**).

**Fig. 4.**
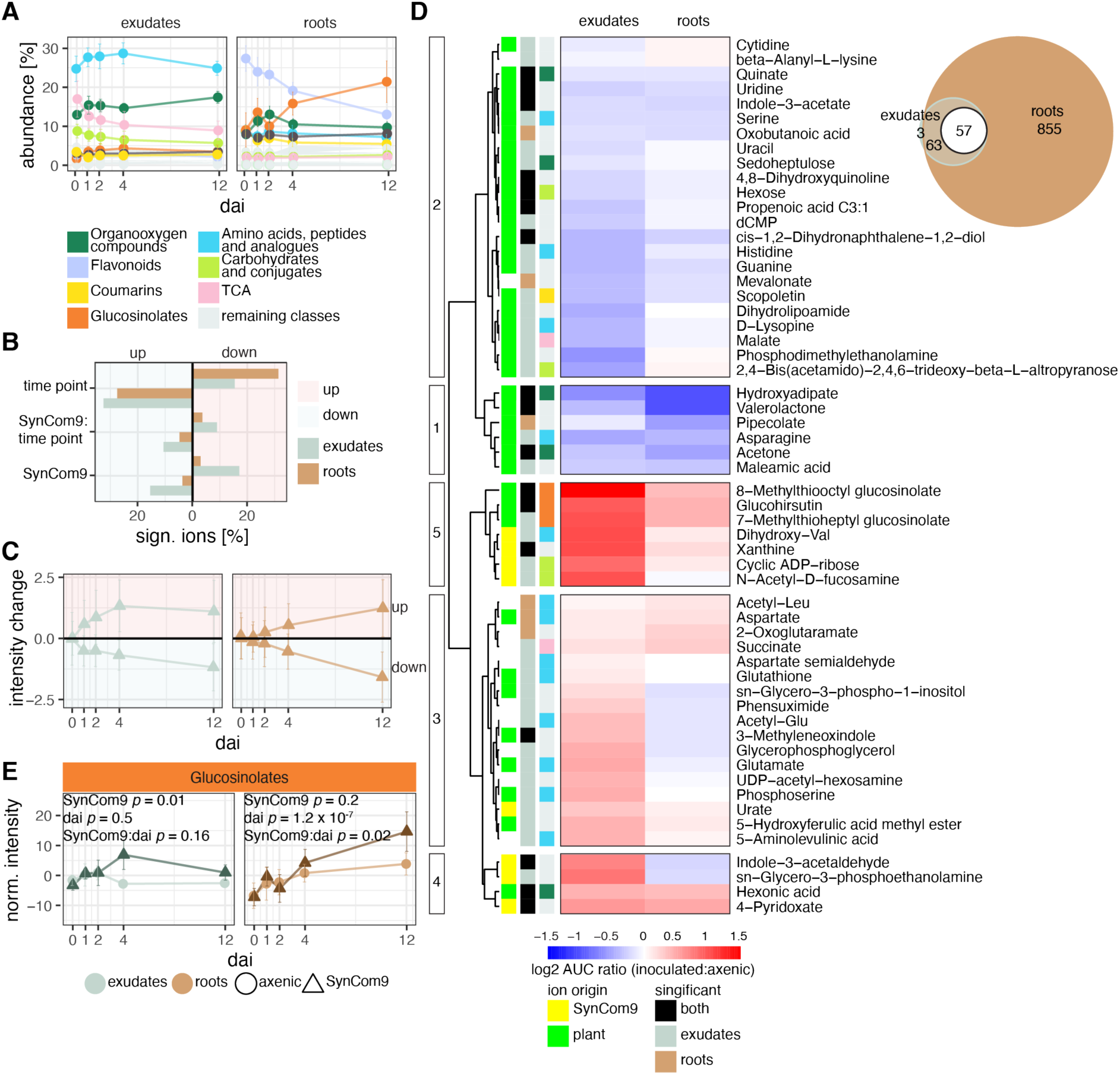
SynCom9-induced metabolic changes followed similar dynamics in roots and exudates. Untargeted analysis (FIA MS) of Arabidopsis exudate and root metabolome dynamics over 12 days in glass-microcosms. Samples were taken 0, 1, 2, 4 and 12 days after inoculation (dai) with the SynCom9. **A** Mean relative intensities of the top-most abundant ion classes (ClassyFire) in exudates and roots over time, with standard deviation (>5% mean relative intensity in at least one sample type). **B** Percentage of ions significantly increased (up: red) or decreased (down: blue) in exudates (blue bars) and roots (brown bars) after FDR *p*-value correction according to two-way ANOVA assessing the effects of time, inoculation, and their interaction. **C** Mean normalised intensity change over time of all ions that were significantly increased (up: red) or decreased (down: blue) in exudates (blue) and roots (brown) in response to inoculation relative to axenic conditions. **D** Hierarchical clustering (complete linkage) and heatmap of the log_2_FC of the area under the curve (AUC) of mean normalised metabolite intensities over time between inoculated and axenic conditions (red: higher, blue: lower in inoculated conditions) in exudates (left) and roots (right). The metabolites depicted were shared between root and exudate data sets and significantly affected by inoculation in at least one sample type (57 metabolites, see Euler plot). Cluster assignments are indicated on the left of the tree. Metabolites are annotated on the left of the heatmap; from left to right *ion origin* (factor estimating the origin of metabolites in exudates of inoculated plants (see *Materials and Methods*), yellow: likely SynCom9-, green: likely plant-derived), *significant* (indicates if metabolites were significantly affected by inoculation in exudates: blue, roots: brown or both: black), metabolite assignment to classes depicted in (**A**). **E** Mean normalised summed metabolite intensities and standard deviations of all metabolites classified as glucosinolates (ClassyFire, **Tables S2, S3**) in exudates (blue, left) and roots (brown, right) over time. Significant changes were determined by two-way ANVOA with Tukey’s HSD and FDR *p*-value correction (N = 8 microcosms per time point à 3 plants each, exudates 123 filtered metabolites, root 975 filtered metabolites).

Most relevant for this work, inoculation with SynCom9 significantly impacted the exudate and root profiles (10% and 3.5% variation, respectively; **Fig. S9, S10)**, but had no significant effect on shoots (**Fig. S11**). To identify metabolites influenced by SynCom9 inoculation in exudates and roots, we compared normalised metabolite intensities over time between inoculated and axenic conditions. Exudates showed a greater proportion of metabolites significantly affected by inoculation (**Fig. 4B**) with more pronounced average intensity changes relative to axenic conditions than roots, particularly at early time points (**Fig. 4C**). However, while the strength of root tissue metabolite responses correlated with the continuous increase of the bacterial load over time, intensity changes in exudates initially increased but levelled off towards later time points (**Fig. S13**). These results indicate that exudate responses to inoculation are stronger but more transient compared to root responses, which are closely linked to bacterial colonization levels.

To understand if changes in specific metabolites followed the same trend in exudates and roots, we compared their changes over time relative to axenic controls for shared metabolites significantly affected by inoculation in at least one sample type (57 metabolites, **Fig. 4D**). Most metabolites showed consistent patterns across exudates and roots. Hierarchical clustering revealed two chemically diverse clusters reduced in both exudates and roots (clusters 1 and 2), two clusters containing several amino acids increasing in exudates (clusters 3 and 4), and, one small cluster containing several aliphatic glucosinolates that increased in both exudates and roots (cluster 5). Glucosinolates stood out as the only major chemical class that grouped in hierarchical clustering and also showed a significant inoculation effect in both exudates and roots (**Fig. 4E**). Several metabolites in clusters 3, 4 and 5 that increased in exudates but not in roots in response to inoculation were likely of bacterial origin based on their abundance in inoculated microcosms in the absence of plants (8 out of the 57 metabolites, **Fig. 4D**). These included N-acetyl-D-fucosamine, a component of bacterial cell walls as well as the bacterial auxin precursor indole-3-acetaldehyde (Etesami and Glick, 2024). Hence, the colonisation of roots by the SynCom9 triggered coordinated responses in both exudates and roots, most notably an increase in glucosinolate levels, and they were accompanied by increased levels of bacterial markers in exudates.

### Exudate-SynCom9 dynamics reveal general and strain-specific patterns

As a final aim of this study, we linked the simultaneously determined exudate and microbiome data to investigate whether variations of specific metabolites might explain load or abundance changes of SynCom9 strains. Hierarchical clustering of individual Pearson correlation coefficients between all SynCom9 strain abundances and all inoculation- responsive metabolite intensities revealed largely consistent patterns across the root-colonizing strains (**Fig. 5A**). Except for *Domibacillus* F004 (barely detected in glass-microcosms, **Fig. 1D**), most strains followed the differential abundance of exudates in response to inoculation. For example, bacterial loads correlated positively with metabolites that were enriched in exudates upon inoculation and were classified as bacteria-derived (**Fig. 5A, B**, clusters 1, 5 and 6). Or, bacterial loads correlated negatively with plant-derived metabolites that were depleted upon colonisation (**Fig. 5A, B**, clusters 2, 3 and 4; e.g. the coumarin scopoletin). However, other metabolites increased upon inoculation but correlated negatively with the bacterial load (e.g. glutamate) or decreased and correlated positively with the bacterial load (e.g. maleamic acid). While the underlying mechanisms leading to these patterns have to be further investigated, they highlight the dynamic interplay between exudation and microbial activity.

**Fig. 5.**
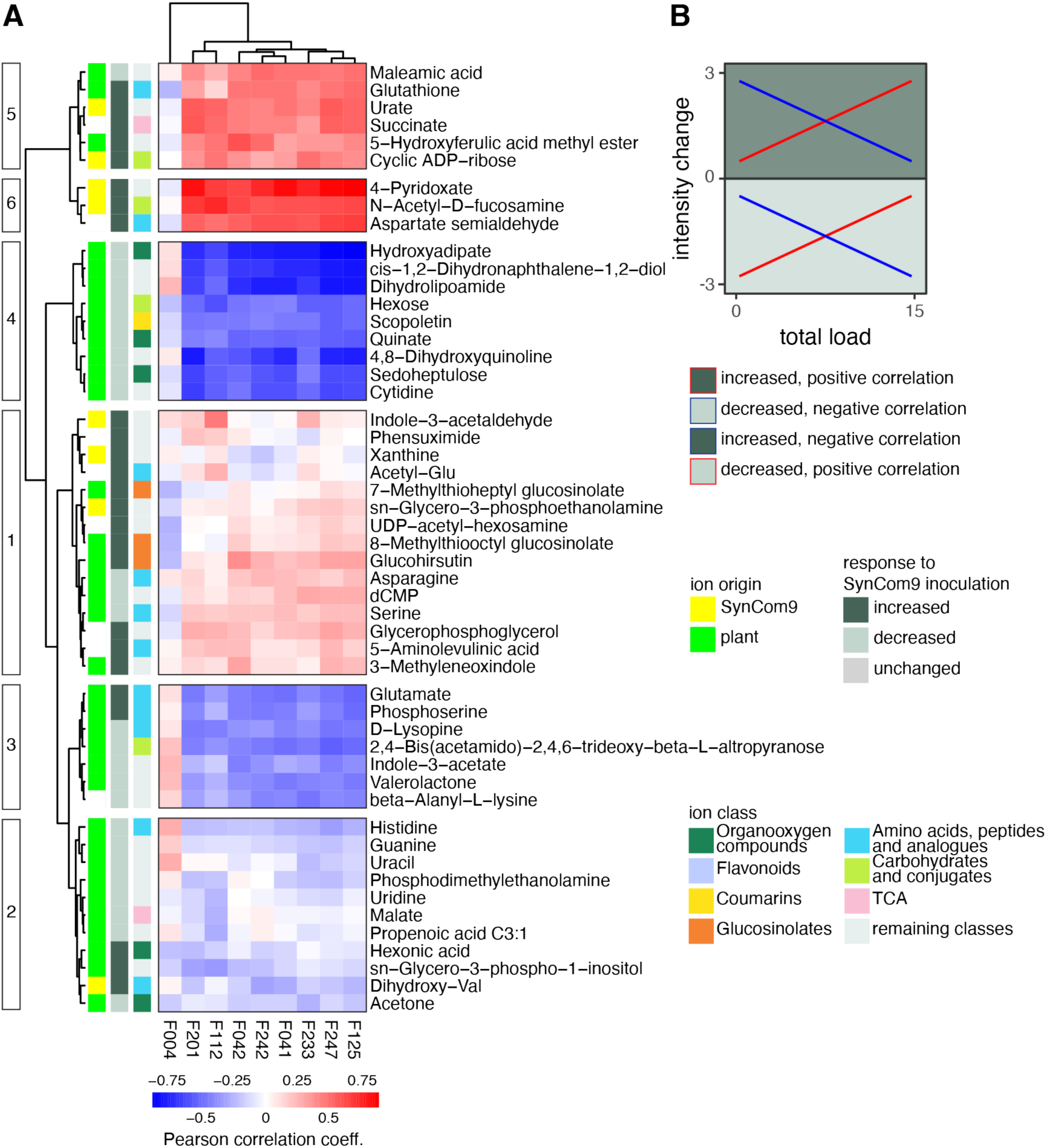
Exudate intensity changes in response to inoculation cannot be linked to the load of single strains. **A** Hierarchical clustering of Pearson correlation coefficients (red: positive correlation, blue: negative correlation) of bacterial loads and root and exudate metabolite changes in inoculated respective to axenic conditions. Only metabolites with significant responses to SynCom9 inoculation are displayed (51 metabolites). Cluster-assignments are indicated on the left of the tree. Metabolites are annotated on the left of the heatmap; from left to right *ion origin* (factor estimating the origin of metabolites in exudates of inoculated plants (see *Materials and Methods*), yellow: likely SynCom9-, green: likely plant-derived), *response to SynCom9 inoculation* (indicates if metabolites were increased (teal) or decreased (blue) in response to inoculation), metabolite assignment to classes depicted in Fig. 4A. Significant changes were determined by two-way ANVOA with Tukey’s HSD and FDR *p*-value correction (N = 40 microcosms à 3 plants each). **B** Explanatory diagram showing the different categories of metabolites in (**A**).

*Pseudomonas* F112, which differed from other strains in its early peak, was no exception to the general pattern. Nevertheless, subtle differences indicate this strain’s particular behaviour. For instance, its load correlated best with the increase of indole-3-acetaldehyde in response to inoculation (**Fig. S14**). Or, in contrast to the general tendency to correlate positively with glucosinolates, *Pseudomonas* F112 showed either no or a negative correlation, suggesting a lower tolerance to glucosinolates. In support of this hypothesis, *Pseudomonas* F112 does not code for homologues of SaxA (while *Pseudomonas* F247 does), a class B beta-lactamase, which is sufficient for the detoxification of aliphatic glucosinolate breakdown products in *Pseudomonas* (Fan et al., 2011, **Table S4**). Thus, this approach uncovered not only overarching patterns but may also be used to identify strain-specific relationships with exudate metabolites.

## Discussion

Disentangling the dynamic metabolic exchange between plants and their associated microbial communities is key to understanding how complex microbiomes are assembled, maintained and how they are adapted over time. However, exudate and microbiome dynamics are seldom studied in parallel, limiting the causal interpretation of dynamic exudate and microbiome relationships. The starting point for this study was a systematic comparison of two microcosm systems: a soil-like clay-microcosm, used in root microbiome studies, and a semi-hydroponic glass-microcosm, developed to study root exudation. Both systems allow similar plant and bacterial growth. Both systems were suitable for most but not all bacteria. The absence of *Domibacillus* F004 from SynCom9 in the glass-microcosms illustrates the limitation of semi-hydroponic conditions to sustain all root-derived bacteria. Not both systems were suitable for exudate measurements: we could not recover exudate signals from clay-grown plants. This highlighted exudate-recovery as a main limiting factor for clay systems, as found in previous studies (Kuijken et al., 2015; Sasse et al., 2020; Strehmel et al., 2014). Our comparison revealed that the semi-hydroponic glass-microcosms are suitable for the paralleled analysis of root exudate and root microbiome dynamics. Therefore, we envisage that this simple and low-cost setup, which allows reproducible and time-resolved measurements, holds potential for advancing the study of exudate–microbe dynamics.

### Microbial dysbiosis and plant performance in microcosm environments

SynCom inoculation did not lead to altered plant growth in both tested growth matrices. This is notable as high-humidity growth conditions in closed systems favour microbial invasions and can lead to a switch of commensal to pathogenic microbial behaviour (Hu et al., 2022; Xin et al., 2016). Here, symptoms such as water-soaking of leaves, associated with high humidity and microbial pathogenicity, were not observed. These findings support the suitability of both microcosm systems for studying plant–microbe interactions under controlled conditions, without confounding effects on overall plant health and development. At the same time, the observed reduction in root biomass in glass-compared to clay-microcosms with unchanged shoot biomass indicates that the physicochemical differences in the plant growth matrix influenced root development without impacting above-ground plant performance. Differences in root biomass may reflect differences in nutrient accessibility between the two growth matrices, for example, due to nutrient absorption by clay particles (López-Bucio et al., 2003).

### Do semi-hydroponic conditions sustain root-associated bacteria?

The limitation of semi-hydroponic conditions for the cultivation of root-derived bacteria was illustrated by the absence of *Domibacillus* F004 in glass bead microcosms. Although in comparison to classical hydroponic systems, glass bead microcosms present more surfaces for bacteria to attach to and a humid but not fully submerged environment, reduced survival of some bacterial taxa was nonetheless to be expected (Stegelmeier et al., 2022; Thomas et al., 2024). However, because *Domibacillus* F004 was not enriched in the root compartment of plants grown in either system, this indicates that this strain is rather clay-than root-associated. However, further studies are needed to identify which root-associated taxa are negatively affected by cultivation with semi-hydroponically-grown plants.

Although root communities in glass-microcosms were initially dominated by *Pseudomonas* strains, over time, low-abundant strains could also establish at higher abundances. Microbiome maturation towards more even and less variable communities has also been observed in soil systems and under field conditions (Almario et al., 2022; Beilsmith et al., 2021; Maignien et al., 2014). So, despite the fact that semi-hydroponic conditions offer limited niches for less competitive strains to persist (Thomas et al., 2024), which was apparent from only few strains recovered from substrate samples, root microbiome assembly seems to follow reproducible and ecologically meaningful trajectories under these simplified and reductionist conditions.

### Exudate effect on microbial communities

In both microcosm types, we observed an exudate effect, marked by a few dominant taxa in the substrate matrix and more even bacterial communities with increased influence of plant roots. This pattern may appear as an “inverse” rhizosphere effect. A rhizosphere effect is typically seen in soil environments, where microbial diversity decreases with proximity to the roots (Bulgarelli et al., 2012; López et al., 2023; Lundberg et al., 2012; Poppeliers et al., 2024; Schlaeppi et al., 2014; Thiergart et al., 2020). It is explained by a selection of bacteria capable of tolerating plant-host specific antimicrobial compounds and more efficiently utilizing plant-derived carbon sources (Basak et al., 2024; Bulgarelli et al., 2013; Thoenen et al., 2023). In this study, however, root-associated bacteria adapted to Arabidopsis were introduced in a carbon-free system where only the Arabidopsis exudates served as a carbon source. Thus, better bacterial performance was expected in the presence of plant roots. In conclusion, in this experimental setup, the plant’s exudates specifically sustain bacterial diversity as there are no alternative carbon sources available.

### Limitations in measuring root exudation in clay-microcosms

Clay-microcosms did not permit the recovery of exudates, especially not *in situ*. Previously, absorption of ca. 20% of exudate compounds of *Brachypodium distachyon* to a clay matrix has been reported (Sasse et al., 2020). Other soil-like matrices with a lower ion-exchange capacity – such as sand, which has been used for exudate sampling of *B. distachyon* or *Lotus japonicus,* for example (Salomonsen et al., 2024; Sasse et al., 2020) – could also be used to enhance *in situ* exudate collection. Sand substrates, though, come with other drawbacks, such as these substrates typically yield lower metabolite recovery than hydroponic systems, and the small particle sizes have been shown to affect root morphology (McLaughlin et al., 2025; Salomonsen et al., 2024; Sasse et al., 2020) and possibly also affect microbial survival.

The more surprising finding here was that the *ex situ* recovery of exudates was diminished from plants previously grown in clay-microcosms. It is conceivable that, although roots were washed for *ex situ* root exudate collection, fine clay particles may remain stuck on the roots, especially on plants grown in unwashed clay, and that these particles absorb the root exudate signal.

The *ex* vs. *in situ* comparison revealed another important finding: plant transfer for *ex situ* root exudate collection results in vastly different exudate profiles compared to *in situ* collected exudates. This might arise from unspecific leakage from possibly wounded root tissue upon transfer, a well-recognised issue in the field that can be (partially) solved by introducing recovery periods before exudate collection (McLaughlin et al., 2023b; Oburger and Jones, 2018; Salomonsen et al., 2024). However, this hardly agrees with time-resolved protocols. Overall, we confirm the non-suitability of clay-microcosms for exudate profiling, both *in* and *ex situ*, and the latter comparison underscored the superior quality of exudate data of plants that do not need to be handled for exudate collection.

### Reproducibility of exudate studies

The two datasets of exudates collected in glass-microcosms (**Fig. S8**), as well as the Arabidopsis exudate profiles measured previously in the same system (Joller et al., 2024) were highly consistent with high relative contributions of organic acids, benzenoids, organoheterocyclic and organooxygen compounds. Further, these exudate profiles are also consistent with the broad exudate profiles in other untargeted exudate analysis (Badri et al., 2013, 2010; Chaparro et al., 2013; McLaughlin et al., 2023b; Salomonsen et al., 2024; Strehmel et al., 2014). Together, this shows the utility of glass-microcosm systems for generating reproducible and broadly comparable exudate profiles in untargeted metabolomic analyses. That being said, the analytics setup used captured mostly polar compounds and thus some well-characterised, apolar, major *Arabidopsis* metabolites previously reported in exudates upon microbial colonisation, such as camalexin (Koprivova et al., 2023; Strehmel et al., 2017), were not detected. Thus, although our analysis encompassed a broad spectrum of metabolic classes, certain key features of Arabidopsis exudates remained undetected, emphasising to the need of future improvements of untargeted metabolomics.

### Exudate responses to inoculation with the commensal SynCom9

We presented proof-of-concept data that the glass-microcosms can be used to measure dynamic microbiome and exudate processes, using root community establishment as an example. We noticed consistently higher levels of aliphatic glucosinolates in exudate and root tissue profiles following inoculation with the SynCom9. Glucosinolates are a well-known class of secondary metabolies in *Brassicaceae* (Halkier and Gershenzon, 2006). They have well-established roles in defence against herbivores (Textor and Gershenzon, 2009) and microbial pathogens (Clay et al., 2009; Strehmel et al., 2017; Finnegan et al., 2016; Lassowskat et al., 2014), and have also been shown to shape microbial communities in Arabidopsis (Bressan et al., 2009; Russ et al., 2024; Unger et al., 2024). Glucosinolates are consistently recovered from Arabidopsis exudates (Mönchgesang et al., 2016) and are enriched in exudates as part of pattern-triggered immunity (Strehmel et al., 2017). Besides immunity, higher levels of glucosinolates have also been associated with a general non-self response to commensal bacteria (Keppler et al., 2024; Maier et al., 2021) and plant-growth promotion by beneficial *Pseudomonas* strains (Jeon et al., 2021). More recently, Arabidopsis-associated bacteria have been shown to use glucosinolates as a carbon source, further suggesting roles of glucosinolates that go beyond immunity (Unger et al., 2024). Future work investigating how glucosinolate release is fine-tuned depending on the presence of beneficials and pathogens, for example, will help shed more light on their diverse functions in plant-microbe interactions. Here, elevated glucosinolate levels in exudates were found after SynCom9 inoculation peaked at early time points. More generally, differences in exudate profiles between inoculated and axenic conditions levelled off after an initial burst, even though the bacterial load continuously increased. This indicates that either a negative feedback loop in plants attenuated responses to commensals with time, that SynCom9 member(s) suppressed exudation of, or degraded metabolites (Ma et al., 2021; Ordon et al., 2025; Teixeira et al., 2021), that responses were connected to abundances of specific strains with varying abundances over time, and/or a combination thereof. For instance, early elevated glucosinolate abundances upon root colonisation may be linked to the *Pseudomonas* dominance. Indeed, microbe-associated molecular patterns eliciting pattern-triggered immunity of commensal *Pseudomonas* strains typically induce strong plant responses (Colaianni et al., 2021). Moreover, glucosinolate detoxifying genes are particularly prevalent in this taxonomic group and are a main determinant of Arabidopsis colonisation success (Keppler et al., 2024; Maier et al., 2021; Russ et al., 2024). Elevated glucosinolate exudation in response to *Pseudomonas* colonisation, paired with the limited capacity of *Pseudomonas* F112 to detoxify glucosinolate-derived isothiocyanates due to the lack of SaxA could thus explain this strain’s drop in abundance over time.

Cases where bacterial degradation of exudates may have contributed to metabolite dynamics not only concern metabolites that were reduced in exudates upon colonisation, but also those that were induced but correlated negatively with the bacterial load. This is particularly interesting in the case of glutamate. Utilisation of dicarboxylates, including glutamate, as a preferred carbon source is a widespread trait among soil-dwelling and plant-associated bacteria, *Pseudomonadaceae* in particular (Lundgren et al., 2021; Yurgel and Kahn, 2004). Recently, it has also been shown that glutamine serves as a chemotactic signal for root colonisers and that the ability to use glutamine as a carbon source contributes to colonisation success (Tsai et al., 2025). It is thus compelling to speculate that root-associated commensal bacteria induce the exudation of preferred carbon sources (Phillips et al., 2004). This could be explored through targeted experiments measuring exudation rates of specific microbial carbon sources in the presence of commensal microbes or their secreted proteins and metabolites.

Analysing exudate profiles and bacterial community composition and loads in parallel allowed us to partially decipher the interplay between bacterial colonization and root exudation, where plant defence responses and bacterial metabolism shape microbial community structure and exudate composition. In future studies, examining the effects of individual bacterial strains and varying inoculation densities on exudate profiles will help to further unravel strain-specific and load-dependent plant exudate responses to colonisation with commensals.

## Materials and Methods

### Plant growth conditions and microcosm setup

We list differences in setups between experiments in **Table S6**. Generally, Arabidopsis seeds were sterilised for 5 min in 70% (v/v) ethanol followed by 12 min 10% (v/v) bleach. Sterile Arabidopsis Col-0 seeds were sown on 0.5 strength Murashige and Skoog (MS) medium plates including (M0222.0025, Duchefa) or excluding (M0221.0050, Duchefa) vitamins supplied with 1% (w/v) phytoagar (P1003.1000, Duchefa). The MS was adjusted to pH 5.7. Plates were stratified for 3-4 days in the dark at 4 °C and then placed vertically in a plant growth chamber (Percival, CFL Plant Climatics, Wertingen, Germany) set to short days with 10 h light (220 µmol m^-2^ s^-1^) (21 °C), and 14 h darkness (18 °C).

Seedlings were pricked to fresh 0.5 strength MS-Agar plates, including or excluding vitamins (**Table S6**) after one week and grown for two more weeks before being transferred to microcosms. Glass-microcosms were prepared as described (McLaughlin et al., 2023a). Briefly, 150 mL of glass beads (⃠ 5 mm) were added to 850 mL glass jars (5743922, Weck glass and packaging GmbH, Bonn, Germany), autoclaved and amended with 35 mL of liquid 0.5×MS. Circa half the glass bead matrix was submerged in growth medium. Clay-microcosms were prepared analogously using clay particles (Attapulgite ⃠ 0.6 – 3 mm, Oil-Dri® special type III R, Maagtechnik, CH) sieved through a 1 mm sieve to remove small particles. To prepare washed clay, sieved clay particles were rinsed with deionised water and dried at 180 °C. Instead of 35 mL, 70 mL – 100 mL of 0.5×MS was added to clay-microcosms; 70 mL to obtain clay particles fully saturated in liquid and 100 mL to obtain fully saturated clay particles plus a liquid phase similar to glass-microcosms (**Table S6**). Depending on the experiment, three to five plants were added (**Table S6**). Microcosms were sealed with glass lids and micropore tape. Note that air-exchange was enhanced by using micropore tape to introduce a space between the lid and the jar (McLaughlin et al., 2023a). Plants were cultivated in microcosms for 1 – 2 weeks. The sterility of non-inoculated microcosms was routinely tested by spotting 20 µL of substrate solution onto plates with tryptic soy agar (TSA, 22092-500G, Sigma-Aldrich) incubated at 28 °C. For the time course experiment (**Fig. 3 – 5**), tissues of single plants within one microcosm were separated during harvest to use for SynCom9 and metabolite profiling. Otherwise, plants from one microcosm were pooled at harvest.

Bacterial cultivation conditions and inoculation of microcosms with the SynCom9 SynCom9 strains (Joller et al., in preperation) (**Table S1**) were streaked onto TSA (F041, F042, F112, F125, F201, F233, F242, F247) or Luria/Miller (LB, x968.3, Carl Roth) agar (F004) from glycerol stocks and cultured up to one week at 28 °C. The SynCom9 inoculum for microcosms was prepared as described (McLaughlin et al., 2023a). Briefly, single colonies were picked to prepare dense suspensions in 800µL of 10 mM MgCl_2_. Strain suspensions were washed three times in 10 mM MgCl_2_ and mixed in equal volumes. This mix was adjusted to a final OD_600_ of 0.002 in 0.5 strength MS, which was used as the substrate solution in microcosms. Microcosms were inoculated at the same day or one day after seedling transfer. SynCom9 densities in microcosms were monitored by making serial dilutions of 100 µL substrate solution plated onto TSA and cultivated for 3 days at 28°C to enumerate the CFU mL^-^_1._

### Exudate, root and shoot metabolomics

#### Sample preparation

Exudates, roots and shoots were harvested at the time points indicated in **Table S6**. Exudates were always collected under axenic conditions. *In situ* exudates were harvested by replacing the growth medium with 50 – 70 mL exudate collection solution equimolar to 0.5×MS (20 mM sterile-filtered ammonium acetate) for 2 h. When comparing root exudate collection in glass- and clay-microcosms, in each microcosm type, enough exudate collection solution was added to fully submerge the growth matrix (50 mL in glass microcosms and 50 – 70 mL in clay-microcosms (depending on absorption by clay). Otherwise, 30 mL of exudate collection solution was added to glass-microcosms. The same procedure was followed with unplanted microcosms to obtain experimental blank samples. For *ex situ* exudate harvesting, plants were retrieved from microcosms and roots were washed by shaking them in exudate collection solution. All plants from a given microcosm were then placed in a Petri dish with the shoots resting on the Petri dish rim containing 50 mL collection solution for 2 h. Petri dishes without plants were prepared as experimental blanks. Exudate samples were centrifuged at 5’000 g for 10 min at 4 °C and supernatants were stored at −80 °C. Before metabolomics analysis, exudates were gently thawed on ice and diluted according to root fresh weights (McLaughlin et al., 2023b).

Frozen roots and shoots were ground to a powder in liquid nitrogen and metabolites were extracted twice on ice in methanol:acetonitrile:ultrapure water 2:2:1 (v/v) for 1 h with intermittent vortexing. After centrifugation at 10’000 g and 4 °C for 5 min, supernatants of both extraction rounds were pooled and dried under vacuum. For metabolite analysis, dried extracts were resuspended in methanol:ultrapure water 1:1 (v/v) according to the tissue fresh weight (200 µL per 70 mg of tissue) and diluted 150 times (McLaughlin et al., 2023b).

#### Metabolomics analysis pipeline

Metabolomics analysis was performed using flow injection-time of flight mass spectrometry (FIA) on a platform consisting of an Agilent 1200 pump coupled to a Gerstel MPS2 autosampler and an Agilent 6550 QTOF mass spectrometer (Agilent, Santa Clara, USA) operating in negative ionisation mode with published parameters (Fuhrer et al., 2011). Briefly, a mobile phase of isopropanol:water 60:40% (v/v) buffered with 5 mM ammonium fluoride was supplied at a rate of 150 µL min^-1^. Homoserine and Hexakis (1H, 1H, 3H-tetrafluoropropoxy)-phosphazine were added to the mobile phase for online mass axis correction. Mass spectra ranging within 50-1000 m/z were recorded at a frequency of 1.4 spectra s^-1^ using maximum resolving power. The source temperature was set to 325 °C, with 5 L min^-1^ drying gas and a nebulizer pressure of 30 spig. Fragmentor, skimmer, and octupole voltages were set to 175 V, 65 V, and 750 V, respectively. Technical blanks (methanol: ultrapure water 1:1 and 20 mM ammonium acetate), sample pools and a standard mixture of 1 µM amino acids were used as quality controls (QC). Samples were injected twice in random order with blank injections between sample groups (QC, exudates, tissues) to reduce sample carryover. The mean of replicated sample measures was taken for data analysis.

Spectral data were processed and annotated according to previously published methods, using the Kyoto Encyclopedia of Genes and Genomes (KEGG) and the Human Metabolome Database (HMDB) (Fuhrer et al., 2011). Metabolites were classified into chemical classes with ClassyFire (Djoumbou Feunang et al., 2016) based on metabolite Kegg IDs converted into international chemical identifiers (InChi). The class of tricarboxylic acid cycle (TCA) metabolites was manually curated. It included the metabolites annotated as pyruvate, fumarate, succinate, oxaloacetic acid, malate, oxoglutarate, aconitate, (iso)citrate, (R)- (homo)3-citrate.

Analysis of FIA data was carried out in R v4.4.2 (R Core Team, 2024). To remove metabolite background noise, FIA data were filtered against experimental blanks (blank glass-microcosms and blank Petri dishes for exudates), technical blanks (methanol:ultrapure water 1:1 for tissues). Metabolites were excluded as background if their median intensities in exudate samples were not at least twice as high as the median intensities in blanks. Sample intensities were normalised using Probabilistic Quotient Normalisation (PQN) (Xia and Wishart, 2011) and for multivariate analysis, metabolites were scaled using autoscaling (van den Berg et al., 2006).

#### Statistical analysis

Analysis was performed in R v4.4.2 (R Core Team, 2024). Differences between sample groups were visualised with Principal Component Analysis (PCA). To quantify effects of experimental factors on metabolite composition, we performed Permutational Multivariate Analysis of Variance (PERMANOVA) with 999 permutations and Euclidean distances using the adonis function from the vegan R package v.2.6.4 (Oksanen et al., 2025). To identify metabolites significantly differing between *in* and *ex situ* glass-microcosm exudate collections, exudate intensities were compared using paired two-sided Student’s t-tests and False Discovery Rate (FDR) *p-*value correction for multiple testing. Else, statistically distinct metabolites were determined using Analysis of Variance (ANOVA) with Tukey’s HSD (honestly significant difference) for post-hoc analysis and FDR *p-*value correction for multiple testing. The origin of metabolites in exudates recovered from glass-microcosms in inoculated conditions was estimated by calculating the log_2_FC ratio between the area under the curve (AUC) of raw metabolite intensities in unplanted, inoculated jars vs planted, axenic jars. A metabolite was assigned as plant or SynCom9 derived if the log_2_FC was at least −0.5 and 0.5, respectively.

### SynCom9 community profiling

#### DNA extraction

For root community profiling, roots were removed from microcosms and glass bead or clay particles sticking to roots were removed. Roots were then tapped dry on paper towels to remove loosely adhering microbes and beat to a slurry with 750 µL of MgCl_2_ 10 mM and sterile glass beads (⃠ 1.7 – 2.1 mm) in a Silamat® S6 (Ivoclar Vivadent, CH) at 50 – 60 Hz for 2 times 30 s. Slurries were kept on ice to avoid overheating. An aliquot of the root slurry was diluted and plated on TSA for CFU enumeration. The rest was stored at −80 °C until processed. Prior to DNA extraction, the root slurry was freeze-dried and ground to a powder under liquid nitrogen. Substrate solutions were spun down twice at maximum speed for 10 min to pellet microbes. DNA was extracted from root powders and substrate pellets in 96-well plates (see **Table S7** for kit specification of single experiments).

DNA extracts were quantified with the AccuClear® Ultra High Sensitivity dsDNA Quantitation Kit (Biotium, Fermont, USA) and, if needed, samples were diluted to 2 ng µL^-1^ with a Myra Liquid Handler (Bio Molecular Systems, Upper Coomera; Australia). Up to 10 ng DNA were used for library preparation.

#### Library preparation

Libraries were prepared using adapted two-step barcoding protocols. In a first PCR (PCR1), either the bacterial 16S V5 – V7 variable gene region (standard PCR) (Bai et al., 2015; Beckers et al., 2016) (**Fig. 1**), or the 16S V5 – V7 variable region together with the Arabidopsis host specific *GIGANTEA* gene (AT1G22770.1) for host-associated microbiome profiling (hamPCR) (Lundberg et al., 2021; Rensburg et al., 2025) (**Fig. 3, 5**) were amplified. Primers and cycling conditions used for specific experiments for PCR1 are listed in **Table S7**. PCR1 amplicons were purified using the ChargeSwitch® PCR Clean-Up Kit (Invitrogen, Waltham, MA, USA) on a KingFisher™ Flex™ Purification System (ThermoFisher Scientific, Waltham, MA, USA). Unique Access Array barcoding primers (Standard BioTools, CA, USA) were added in a second PCR (PCR2) using 5 µL of purified PCR1 product following previously reported cycling conditions (Rensburg et al., 2025). The number of cycles depended on the library preparation protocol: 10 cycles for standard 16S V5 – V7 amplification and 25 cycles for hamPCR. In total, both library preparation protocols entailed 35 PCR cycles. PCR2 products were purified and quantified as before. 100 ng of each sample was pooled using a Myra Liquid Handler to create libraries. Libraries were paired-end sequenced on an Illumina MiSeq instrument at the NGS platform of the University of Bern (https://www.ngs.unibe.ch) with the chemistries listed in **Table S7**. Given the low diversity of the SynCom9, shallow sequencing resulting in an average of 4228 reads sample^-1^ was sufficient to cover the diversity of our samples (**Fig. S15**).

#### Bioinformatics

MiSeq reads were quality checked with FastQC v0.11.8 (Babraham Institute, Cambridge, United Kingdom). The NGS platform removed the barcodes from sequences and wrote them to the sequence headers. The data were demultiplexed and primers were removed from sequences using cutadapt v3.4 (Martin, 2011). Quality filtering, read merging and amplicon sequence variant (ASV) inferring were performed as described (Gfeller et al., 2023) with the dada2 package v1.26.0 (Oksanen et al., 2024) in R v4.2.0 (R Core Team, 2022). Taxanomic assignments were done with a naïve Bayesian classifier in a DADA2 formatted version from the SILVA v. 132 database (Quast et al., 2013). Taxonomy assignments were refined with the IDTAXA classifier using the SILVA r138 2019 via DECIPHER v2.26.0 (Wright, 2016) and replaced wherever IDTAXA assigned more taxonomic levels than the naïve Bayesian classifier. In the library prepared with hamPCR, ASVs assigned as Eukarya were aligned with the *GIGANTEA* sequence (AGAGTATGGAGCTGGGATTGACTCGGCAATTAGTCATACGCGCCGAATTTTGGCAATCCTAGAGGCCTCTTTTCATTAAAACCATCTTCTGTGGGGACTCCATGGAGTTACAGTTCTAGTGAGATAGTTGCTGCGGCCATGGTTGCAGCTCATATTTCCGAACTGTTCAGACGTTCAAAGGCCTTGACGCATGCATTGTCTGGGTTGATGAGATGTAAGTGGGATAAGGAAATTCATAAAAGAGCATCATCATTATATAACCTCATAGATGTTCACAGCAAAGTTGTTGCCTCCA) and kept if the sequence similarity was > 99%, which yielded a single *GIGANTEA* host ASV. Remaining ASVs annotated as Eukarya, as well as Archaea, Cyanobacteria or Mitochondria were removed. ASVs were assigned to SynCom9 strains if their sequence corresponded 100% to a 16S V5 – V7 gene amplicon sequence of a SynCom9 strain. SynCom9 amplicon sequences were obtained from libraries as prepared here, sequenced on an Illumina MiSeq instrument (v2 chemistry, 500 bp paired end). Computations were carried out at the sciCORE (http://scicore.unibas.ch/) scientific computing centre of the University of Basel.

#### Statistical analysis

For statistical analyses, we used the vegan package v2.6.4 (Oksanen et al., 2025) in R v4.4.2 (R Core Team, 2024). Following published recommendations (Weiss et al., 2017), raw reads were normalised by rarefaction. Rarefaction thresholds used for the single libraries are specified in **Table S7**. For the library prepared with hamPCR (**Table S7**), the microbial load of each sample was calculated as described before (Lundberg et al., 2021). We calculated differences between sample groups based on Bray-Curits dissimilarity matrices using rarefied count tables or, for the library prepared with hamPCR, rarefied count tables scaled by the microbial load with the R package phyloseq (McMurdie and Holmes, 2013). Bray-Curtis dissimilarities were visualised with unconstrained Principal Coordinate Analysis (PCoA). To quantify the effects of experimental factors on Bray-Curtis dissimilarities, we performed PERMANOVA with 999 permutations using the adonis function from the vegan package.

### Homology search

SynCom9 genomes were sequenced, assembled and annotated as previously described (Joller et al., in preperation). For homology search, a fasta file containing protein sequences of interest was created: SaxA, SaxB, SaxF, SaxG were copied from the *Pseudomonas syringae* pv. tomato reference genome, SaxC (Q885H6), SaxD (Q87X84, this protein is annotated as SaxB on Uniprot but was originally named SaxD (Fan et al., 2011) and fliC (P21184) were retrieved from Uniprot. With the blast package v.2.15.0, a blast database was created for each genome using the function *makeblastdb* and then homologs were searched in each genome database with *tblastn*. Extracted fliC sequences were aligned with published flg22 sequences to evaluate extracted flg22 sequences and estimate their immunogenicity (Colaianni et al., 2021).

## Data and code availability

The source data and scripts used for the data analysis performed in this study will be made publicly available on GitHub (https://github.com/PMI-Basel/Joller_et_al_Glass_Clay). The raw sequencing data are stored at the European Nucleotide Archive (http://www.ebi.ac.uk/ena) under the study accession PRJEB107228.

## Acknowledgements

We sincerely thank Prof. Dr. Nicola Zamboni (ETH Zurich, Switzerland) for generating the metabolome data. We would also like to acknowledge Dr. Henry Janse van Rensburg for providing support in library preparation, and Drs. Sarah McLaughlin, Alexandra Siffert and Delphine Chinchilla for their assistance in establishing the clay and glass bead microcosm systems. This work was supported by a Plant Science Centre-Syngenta research fellowship awarded to K.S. and J.S., supporting C.J.

## Author Contributions

C.J., K.S. and J.S. designed the experiments and wrote the manuscript. C.J. performed the experiments. J.W. performed bioinformatics analysis of amplicon sequencing data and Sax homology search. C.J. analysed the data. K.S. and J.S. supervised the project.

