## Supplemental Figures and Tables for "Paralleled Dynamics of Arabidopsis Root Exudation and SynCom Assembly in a Controlled Environment"

### Supplementary Figures

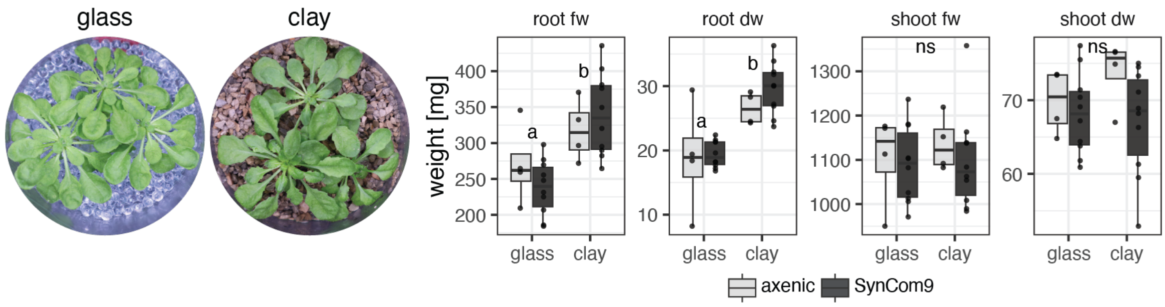


**Fig. S1 Comparable plant biomass in glass- and clay-microcosms.** Summed weights of plants grown in axenic (lightgrey) and SynCom9-inoculated (darkgrey) glass- and clay-microcosm (N = 10). Boxplots show the median with first and third quartiles. The hinges represent the range of data within 1.5 times the interquartile range (IQR). Significantly distinct weights are indicated with letters (ANOVA and Tukey's HSD, ns: not significant).

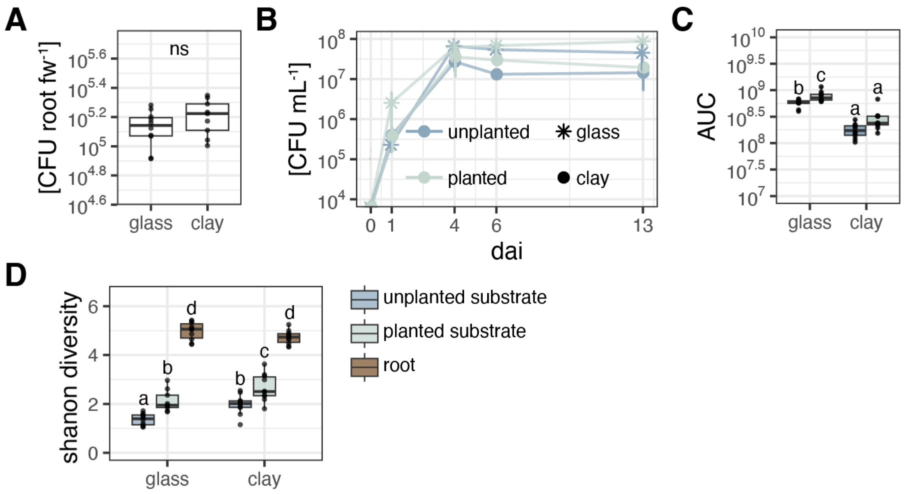


**Fig. S2 An exudate effect in microcosms is measurable in higher microbial diversity with increasing plant influence. A**, **B** Colony forming units (CFU) in the substrate of glass- (asterisk symbols) and clay- (circles) microcosms with (light colour) and without (dark colour) plants. Data is displayed as means with the standard deviation per timepoint (days after inoculation, dai, N = 10). **B** Area under the curve (AUC) of curves displayed in **A**. The data were normalised by mean CFU counts at 0 dai. Letters indicate significant differences according to ANOVA and Tukey's HSD (N = 10). **C** Colony forming units normalised by root fresh weight of plants harvested from glass- and clay-microcosms at 13 dai (N = 10). **D** Shannon diversity index of SynCom9 communities in glass and clay microcosms of unplanted substrate (dark blue), planted substrate (light blue), and on roots (brown). Letters indicate significant differences according to ANOVA and Tukey's HSD (N = 9-10). Boxplots show the median with first and third quartiles. The hinges represent the range of data within 1.5 times the interquartile range (IQR).

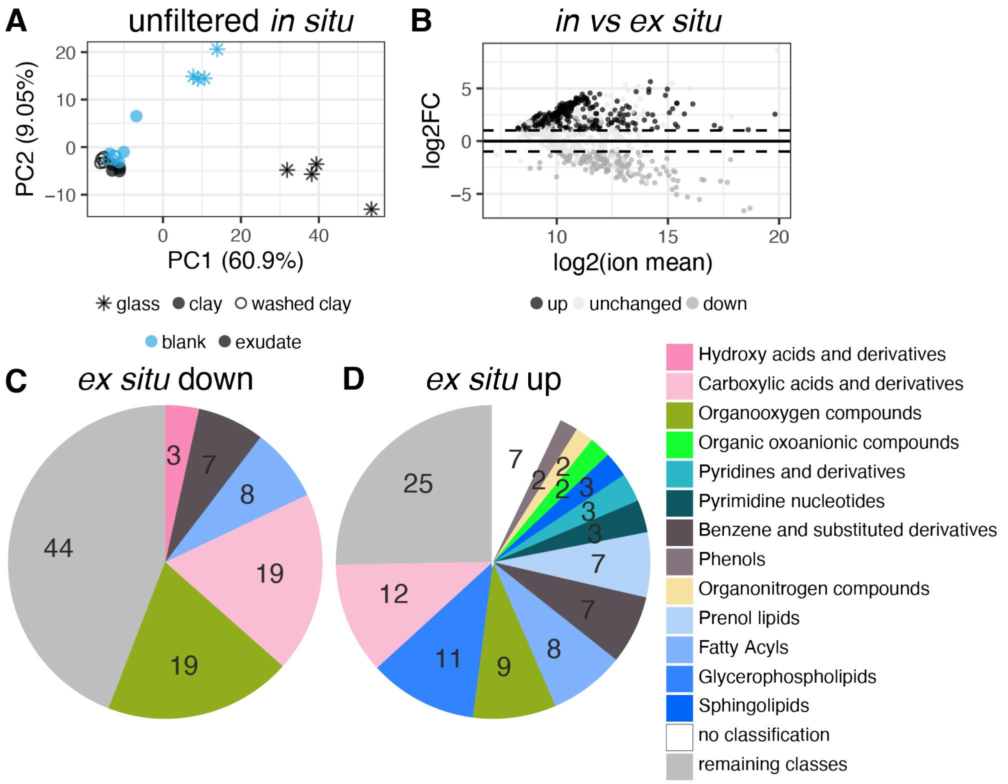


**Fig. S3 Recovered exudates differ depending on plant growth conditions (glass- and clay-microcosms) and exudate collection mode (*in* and *ex situ*). A** Principal Component Analysis (PCA) of normalised intensities of the unfiltered metabolite data set (2135 metabolites). Displayed are exudates (black) and blanks (blue) collected *in situ* from glass- (asterisk symbols), clay- (circles) and washed clay-microcosms (open circles, N = 3-4 microcosms per condition comprising 5 plants each). The filtered data set and *ex situ*-collected exudates are shown in **Fig. 2A**. **B** MA plot displaying $log_2FC$ of exudate intensities collected *in* and *ex situ* from glass-microcosms and $log_2FC$ mean metabolite intensities across both conditions (black: more, grey: less abundant *ex situ* and lightgrey: unchanged). **C, D** Number of metabolites pre class that were down **C** or up **D** in *ex situ*-compared to *in situ*-collected exudates from glass-microcosms. **B-D** Student's t-test with FDR p-value correction for multiple testing ($p \leq 0.05$, N = 4).

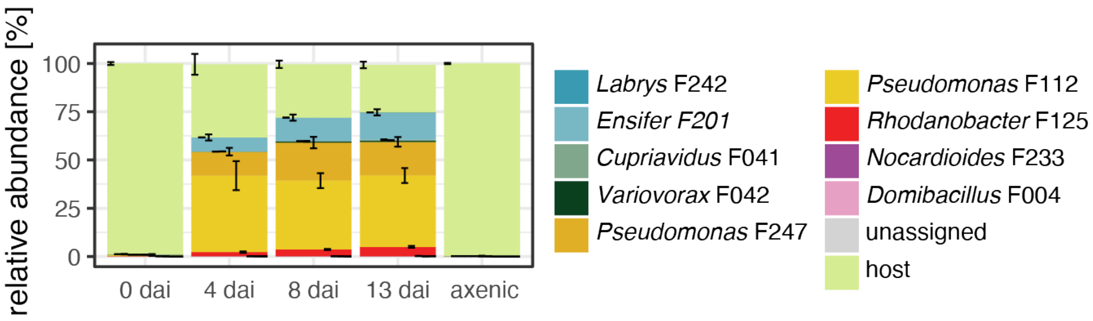


**Fig. S4 Pilot data indicates that SynCom9 root communities establish within 4 days.** Mean relative abundances of SynCom9 and Arabidopsis *GIGANTEA* (host) amplicons in root samples as determined by hamPCR (Lundberg et al., 2021) in a pilot experiment with glass-microcosms and 4 harvesting time points (0, 4, 8 and 13 dai). Axenic: roots harvested from non-inoculated glass microcosms. Displayed are means with standard deviations (N = 8 microcosms à 3 plants each).

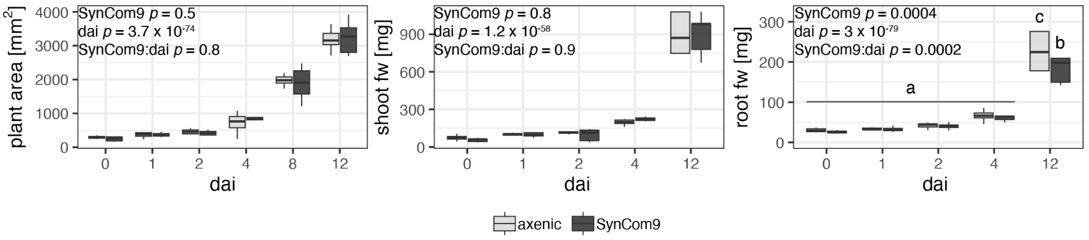


**Fig. S5 No effect of inoculation on shoot, but late effects on root biomass in glass microcosms.** Rosette area, shoot and root fresh weight per microcosm (sum of three plants) inoculated with the SynCom9 (black) or a sterile mock treatment (grey). Letters indicate significant differences according to ANOVA and Tukey's HSD. Post-hoc analysis was only carried out if there was a significant effect of SynCom9 inoculation (dai: harvesting time point, axenic: tissues harvested from non-inoculated microcosms, N = 8 microcosms per time point à 3 plants each). Boxplots show the median with first and third quartiles. The hinges represent the range of data within 1.5 times the interquartile range (IQR).

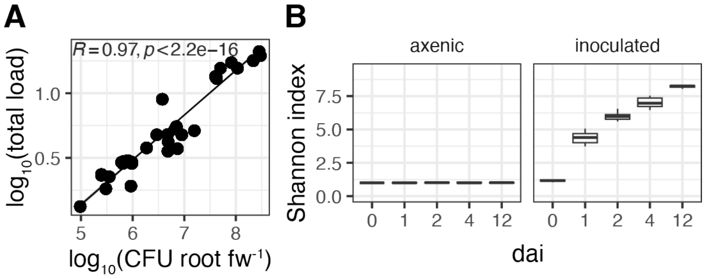


**Fig. S6 The bacterial load correlates well with root CFU numbers and root SynCom9 community diversity increases in glass-microcosms with time. A** Pearson correlation between the $log_{10}$ of the total bacterial load and normalised root CFU. The bacterial load corresponds to the bacteria-to-host amplicon ratio as determined by hamPCR (Lundberg et al., 2021). The grey area shows 95% confidence intervals. The Correlation coefficient and *p*-value are indicated on the plot. **B** Shannon diversity index of root bacterial communities in mock-inoculated (axenic: left panel) and SynCom9-inoculated (inoculated: right panel) microcosms. Boxplots show the median with first and third quartiles. The hinges represent the range of data within 1.5 times the interquartile range (IQR). Tissues were harvested at 5 different time points (0, 1, 2, 4 and 12 dai, N = 8 microcosms per time point à 3 plants each).

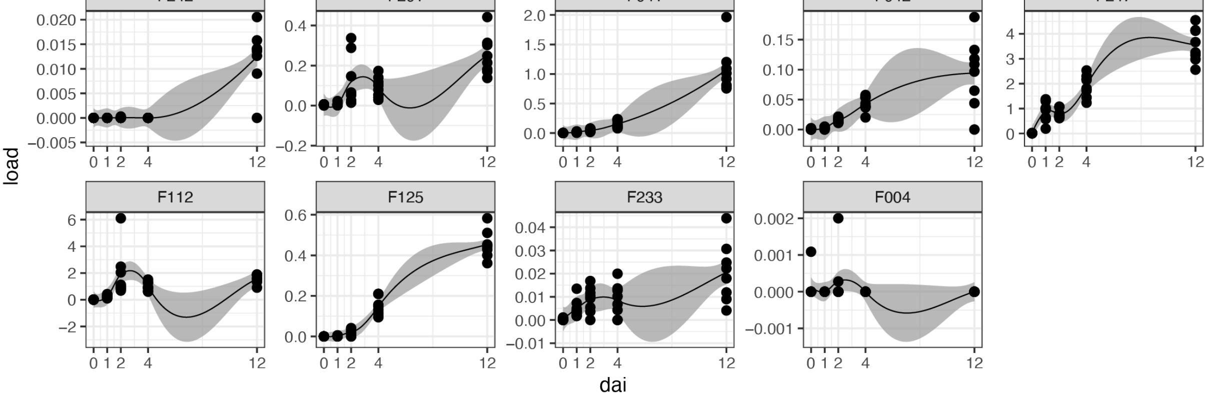


**Fig. S7 Root colonization dynamics in glass microcosms of single strains shows a general trend for increased abundances with time.** Bacterial load of single SynCom9 strains on roots of Arabidopsis plants cultivated in glass microcosms. Singel data points and LOESS regression with 95% confidence intervals are shown. The bacterial load corresponds to the bacteria-to-host amplicon ratio as determined by hamPCR (Lundberg et al., 2021). Tissues were harvested at 5 different time points (0, 1, 2, 4 and 12 dai, N = 8 microcosms per time point à 3 plants each).

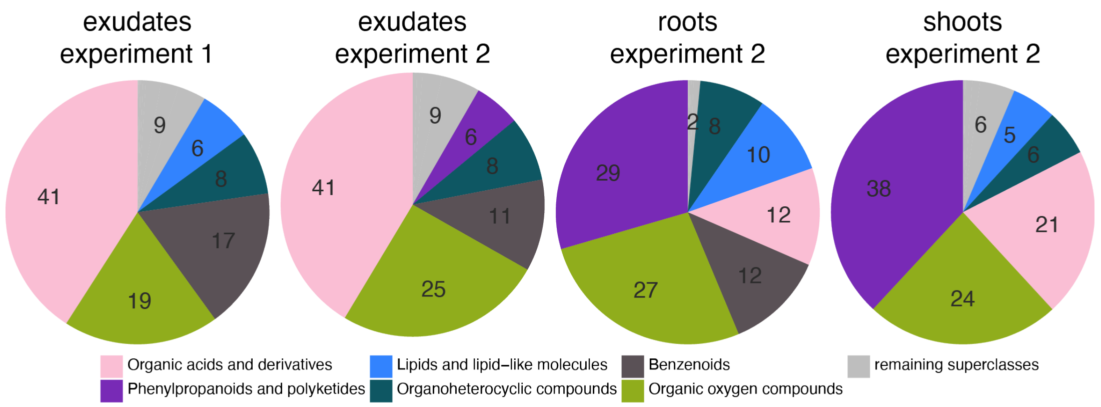


**Fig. S8 The chemical composition of exudates was consistent across experiments but differed from roots and shoots.** Pie charts displaying the ion intensity proportion of ClassyFire metabolite superclasses averaged across all samples in exudates (715 and 123 metabolites in experiment 1 (**Fig. 2**) and 2 (**Fig. 4**), respectively), roots (975 metabolites) and shoots (838 metabolites).

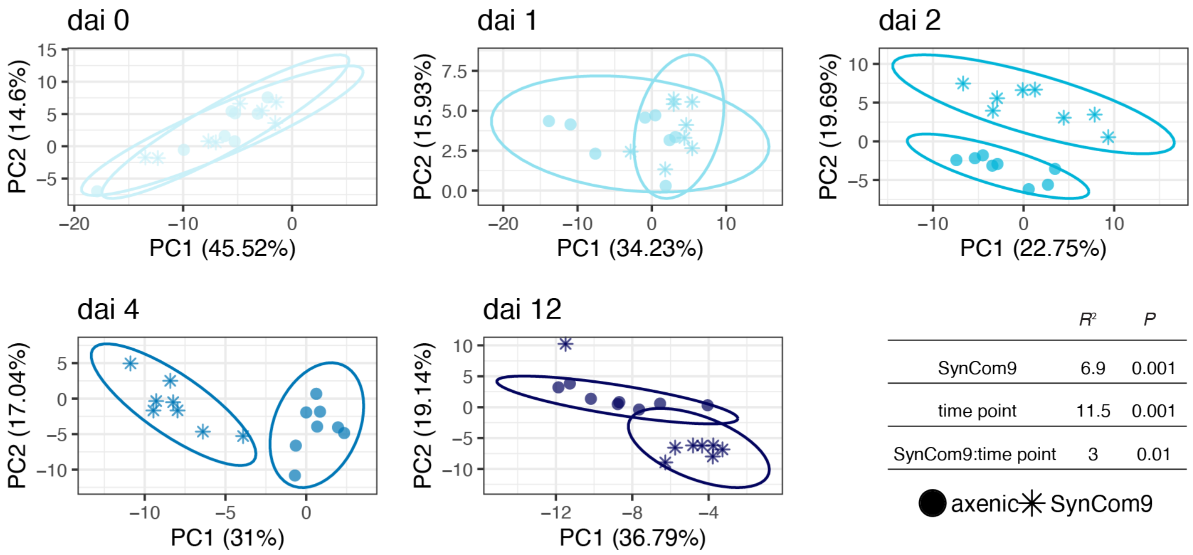


**Fig. S9 Exudate responses to inoculation over time in glass microcosms.** Principal component analysis of normalised exudate metabolite intensities in sterile versus inoculated samples. Ellipses represent 95% confidence intervals. The table shows PERMANOVA results calculated using Euclidean distances (123 filtered metabolites, N = 8 microcosms per time point à 3 plants each).

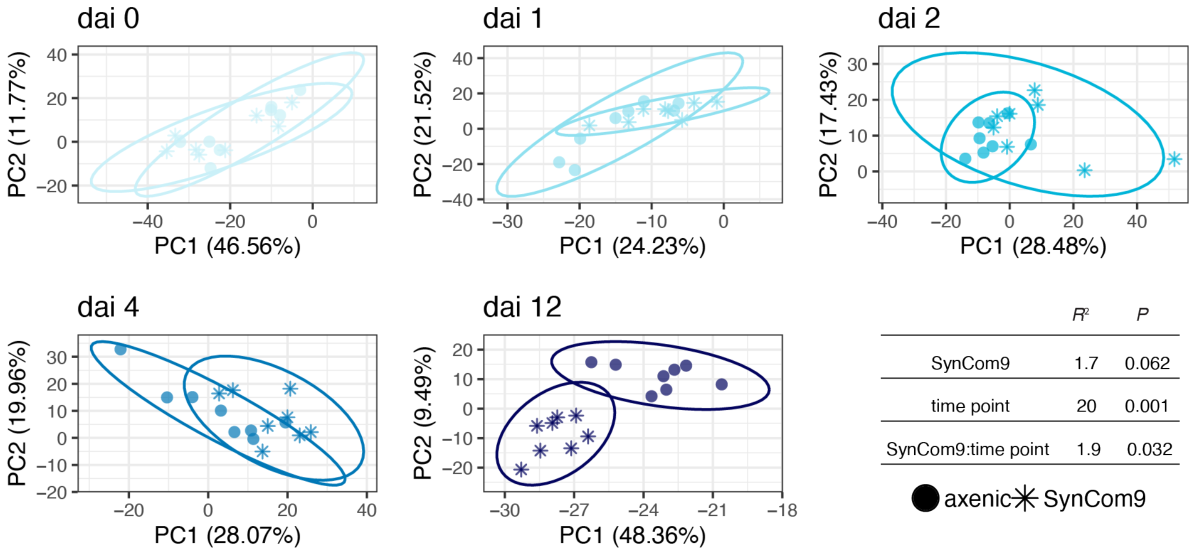


| | $R^2$ | $P$ |
|---|---|---|
| SynCom9 | 1.7 | 0.062 |
| time point | 20 | 0.001 |
| SynCom9:time point | 1.9 | 0.032 |

**Fig. S10 Root responses to inoculation over time in glass microcosms.** Principal component analysis of normalised root metabolite intensities in sterile versus inoculated samples. Ellipses represent 95% confidence intervals. The table shows PERMANOVA results calculated using Euclidean distances (975 filtered metabolites, N = 8 microcosms per time point à 3 plants each).

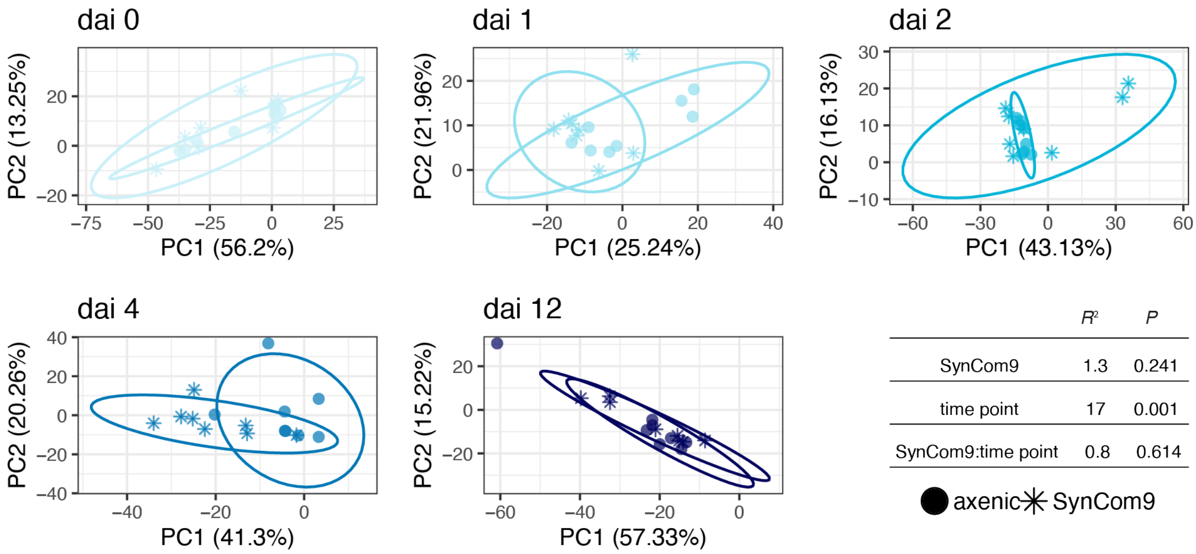


| | $R^2$ | $P$ |
|---|---|---|
| SynCom9 | 1.3 | 0.241 |
| time point | 17 | 0.001 |
| SynCom9:time point | 0.8 | 0.614 |

**Fig. S11 Shoot responses to inoculation over time in glass microcosms.** Principal component analysis of normalised shoot metabolite intensities in sterile versus inoculated samples. Ellipses represent 95% confidence intervals. The table shows PERMANOVA results calculated using Euclidean distances (838 filtered metabolites, N = 8 microcosms per time point à 3 plants each).

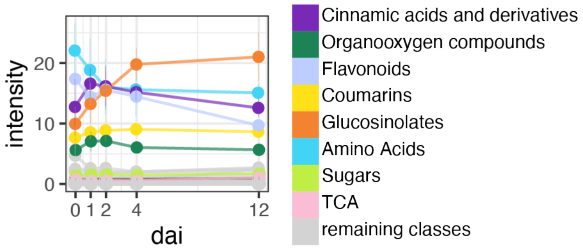


**Fig. S12 Dynamic change of shoot metabolite classes over time.** Mean shoot metabolite class intensities with standard deviation over time averaged across samples are displayed for the top most abundant classes (>5% mean intensity in exudates, roots or shoots, 838 filtered metabolites, N = 8 microcosms à 3 plants each, TCA: tricarboxylic acids).

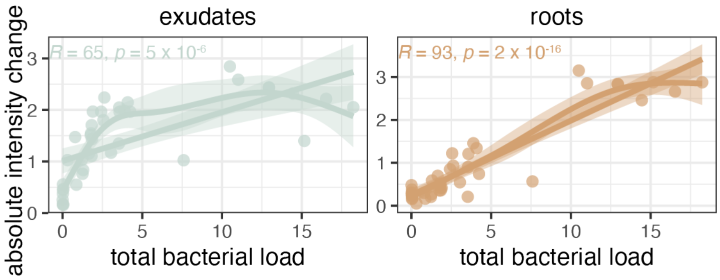


**Fig. S13 Linear correlation between absolute intensity changes in response to inoculation and the total bacterial load is better for roots than exudates.** Pearson correlation of the total bacterial load as determined by hamPCR (Lundberg et al., 2021) with absolute mean normalised intensity changes of all metabolites that were significantly affect by SynCom9 inoculation relative to axenic conditions in exudates (left panel, blue) and roots (right panel, brown). Significant metabolites were determined by two-way ANVOA with Tukey's HSD and FDR *p*-value correction. Pearson correlation coefficients and *p*-values as well as best model fits according to a Generalised Additive Model (GAM) are indicated in the plot. The shaded areas show 95% confidence intervals (N = 40 microcosms à 3 plants each).

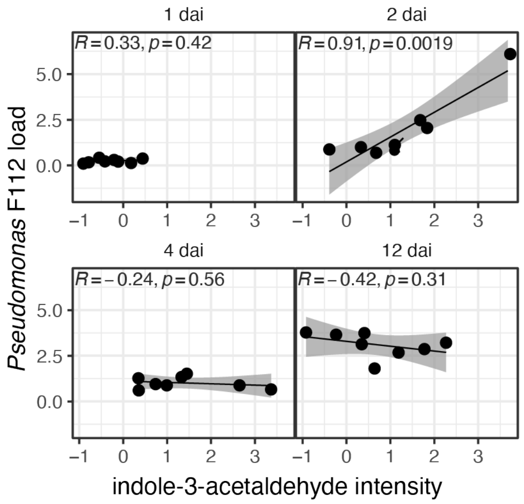


**Fig. S14 The *Pseudomonas* F112 load on roots correlated well with indole-3-acetaldehyde intensities in exudates at the time point where it was dominant in root SynCom9 communities.** Pearson correlations of *Pseudomonas* F112 load with indole-3-acetaldehyde intensities for single timepoints. Correlation coefficients and *p*-values are indicated on the top left. 95% confidence intervals are indicated as grey areas (N = 8 microcosms à e plants each).

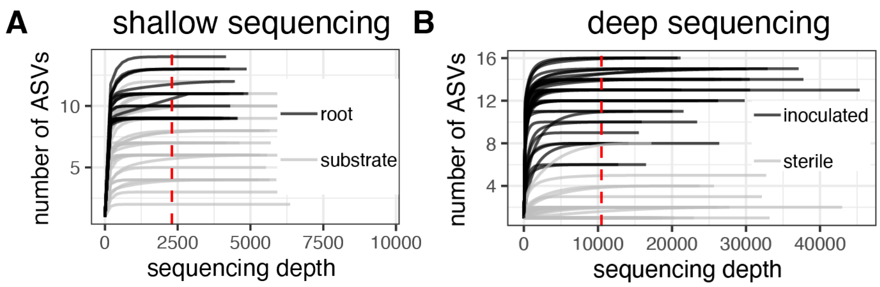


**Fig. S15 Shallow sequencing is sufficient to cover the SynCom9 diversity.** Rarefaction curves of amplicon sequencing samples shown in **A Fig. 1** and **B Fig. 3**. Libraries were sequenced using a **A** shallow- and **B** deep-sequencing approach (**Table S7**). The red dotted line indicates the rarefaction threshold used for normalization. ASV: Amplicon sequencing variant.

### Supplementary Tables

**Table S1 SynCom9 community members.**

| Strain ID | Phylum | Class | Order | Family | Genus |
|---|---|---|---|---|---|
| F201 | *Alphaproteobacteria* | *Alphaproteobacteria* | *Rhizobiales* | *Rhizobiaceae* | *Ensifer* |
| F242 | *Alphaproteobacteria* | *Alphaproteobacteria* | *Rhizobiales* | *Labraceae* | *Labrys* |
| F041 | *Betaproteobacteria* | *Betaproteobacteria* | *Burkholderiales* | *Burkholderiaceae* | *Cupriavidus* |
| F042 | *Betaproteobacteria* | *Betaproteobacteria* | *Burkholderiales* | *Comamonadaceae* | *Variovorax* |
| F112 | *Gammaproteobacteria* | *Gammaproteobacteria* | *Pseudomonadales* | *Pseudomonadaceae* | *Pseudomonas* |
| F125 | *Gammaproteobacteria* | *Gammaproteobacteria* | *Xanthomonadales* | *Rhodanobacteraceae* | *Rhodanobacter* |
| F247 | *Gammaproteobacteria* | *Gammaproteobacteria* | *Pseudomonadales* | *Pseudomonadaceae* | *Pseudomonas* |
| F004 | *Firmicutes* | *Bacilli* | *Bacillales* | *Planococcaceae* | *Domibacillus* |
| F233 | *Actinobacteria* | *Actinobacteria* | *Propionibacteriales* | *Nocardioidaceae* | *Nocardioides* |

**Table S2 Exudate metabolites annotated as glucosionaletes with ClassyFire classifications. Bold compound names indicate metabolites that were significantly affected by inoculation.**

| ionMz | formula | ion | score | TopAnnotationName | KEGGid | Class | Parent Level 3 |
|---|---|---|---|---|---|---|---|
| 420.0465 | C12H23NO9S3 | -H(-) | 100 | Glucoerucin | C08409 | Organooxygen compounds | Glucosinolates |
| 462.0932 | C15H29NO9S3 | -H(-) | 100 | **7-Methylthioheptyl glucosinolate** | C17252 | Organooxygen compounds | Glucosinolates |
| 476.1093 | C16H31NO9S3 | -H(-) | 100 | **8-Methylthiooctyl glucosinolate** | C17254 | Organooxygen compounds | Glucosinolates |
| 477.0647 | C17H22N2O10S2 | -H(-) | 100 | 4-Methoxyglucobrassicin | C08423 | Organooxygen compounds | Glucosinolates |
| 492.1038 | C16H31NO10S3 | -H(-) | 100 | **Glucohirsutin** | C17271 | Organooxygen compounds | Glucosinolates |

**Table S3 Root metabolites annotated as glucosinolates with ClassyFire classifications. Bold compound names indicate metabolites that were significantly affected by inoculation.**

| ionMz | formula | ion | score | TopAnnotationName | KEGGid | Class | Parent Level 3 |
|---|---|---|---|---|---|---|---|
| 420.0465 | C12H23NO9S3 | -H(-) | 100 | Glucoerucin | C08409 | Organooxygen compounds | Glucosinolates |
| 422.0259 | C11H21NO10S3 | -H(-) | 100 | **Glucoiberin** | C08411 | Organooxygen compounds | Glucosinolates |
| 436.0417 | C12H23NO10S3 | -H(-) | 100 | **Glucoraphanin** | C08419 | Organooxygen compounds | Glucosinolates |
| 450.0566 | C13H25NO10S3 | -H(-) | 100 | **Glucoalyssin** | C08400 | Organooxygen compounds | Glucosinolates |
| 452.0354 | C12H23NO11S3 | -H(-) | 100 | **Glucoerysolin** | C08410 | Organooxygen compounds | Glucosinolates |
| 462.0932 | C15H29NO9S3 | -H(-) | 100 | **7-Methylthioheptyl glucosinolate** | C17252 | Organooxygen compounds | Glucosinolates |

| | | | | | | | |
|---|---|---|---|---|---|---|---|
| 476.1093 | C16H31NO9S3 | -H(-) | 100 | 8-Methylthiooctyl glucosinolate | C17254 | Organooxygen compounds | Glucosinolates |
| 477.0647 | C17H22N2O10 S2 | -H(-) | 100 | 4-Methoxyglucobrassicin | C08423 | Organooxygen compounds | Glucosinolates |
| 492.1038 | C16H31NO10S 3 | -H(-) | 100 | **Glucohirsutin** | C17271 | Organooxygen compounds | Glucosinolates |

**Table S4 Number of Sax (Survival on Arabidopsis extracts) homologues per SynCom9 strain.**

| Strain | Genus | SaxA | SaxB | SaxC | SaxD | SaxF | SaxG |
|---|---|---|---|---|---|---|---|
| F004 | *Domibacillus* | 0 | 1 | 0 | 2 | 2 | 3 |
| F041 | *Cupriavidus* | 0 | 3 | 10 | 11 | 10 | 15 |
| F042 | *Variovorax* | **1** | 4 | 8 | 19 | 18 | 22 |
| F112 | *Pseudomonas* | 0 | 3 | 10 | 11 | 10 | 15 |
| F125 | *Rhodanobacter* | 0 | 0 | 1 | 16 | 13 | 16 |
| F201 | *Ensifer* | **1** | 6 | 11 | 11 | 11 | 14 |
| F233 | *Nocardioides* | 0 | 2 | 5 | 0 | 0 | 0 |
| F242 | *Labrys* | 0 | 8 | 11 | 16 | 15 | 18 |
| F247 | *Pseudomonas* | **1** | 5 | 12 | 10 | 10 | 12 |

**Table S5 Immunogenicity of SynCom9 flg22 epitopes as determined by flg22 sequence identity with flg22 epitopes reported by Colaianni and colleagues 2021.** Epitopes with a high ROS-Zscore and a low SGI-Zscore are more immunogenic.

| Epitope_ID | Query Cover | E value | % identity | length | strain | ROS-Zscore | SGI-Zscore_effect | fdr_pvalue | Clade |
|---|---|---|---|---|---|---|---|---|---|
| **Pta** | 100% | 8.00E-21 | 100 | 22 | **F247** | 6.39141625 | -1.872072143 | 1.11E-36 | 1 |
| **1983** | 100% | 4.00E-21 | 100 | 22 | **F125** | 6.221189735 | -1.988952962 | 7.72E-09 | 1 |
| **295** | 100% | 1.00E-18 | 90.91 | 22 | **F004** | 4.393882378 | -1.394847159 | 7.11E-06 | 2 |
| **289** | 100% | 8.00E-21 | 100 | 22 | **F004** | 4.204316273 | -0.934446049 | 0.003248079 | 2 |
| **289** | 100% | 1.00E-18 | 90.91 | 22 | **F004** | 4.204316273 | -0.934446049 | 0.003248079 | 2 |
| **5013** | 95% | 3.00E-20 | 100 | 22 | **F041** | 3.670873321 | -1.311582294 | 0.000308889 | 1 |
| **5013** | 95% | 3.00E-20 | 100 | 22 | **F112** | 3.670873321 | -1.311582294 | 0.000308889 | 1 |
| **94** | 100% | 4.00E-21 | 100 | 22 | **F233** | 3.602624999 | -0.376858768 | 0.334350529 | 2 |
| **1471** | 100% | 2.00E-19 | 95.45 | 22 | **F004** | 3.472763109 | -0.343345359 | 0.312093402 | 2 |
| **1329** | 100% | 1.00E-21 | 100 | 22 | **F042** | 0.053434509 | -0.813765196 | 0.03775908 | 1 |

**Table S6 Overview of microcosm experiments conducted in this study.**

| **Experiment** | **Growth medium** | **Plants per microcosm** | **Day of transfer (dat)** | **Day of inoculation (dai)** | **Sampling (dat)** | **CFU substrate (dai)** | **Sample types** (number of plants per microcosm used for sample type) | **Volume for exudate collection** | **Figures** |
|---|---|---|---|---|---|---|---|---|---|
| Microbiome in glass- vs clay-microcosms | 0.5 MS including vitamins | 3 | 21 | 0 | 14 | 0, 1, 4, 6, 13 | SynCom9 community (3) | no exudates harvested | 1, S1, S2, S15 |
| Exudate sampling in glass- and clay-microcosms | 0.5 MS including vitamins | 5 | 21 | no inoculation | 7 | | Exudates (5) | 50 – 100mL | 2, S3, S8 |
| Microbiome and exudate dynamics in glass-microcosms | 0.5 MS excluding vitamins | 3 | 21 | 1 | 1, 2, 3, 5, 13 | 0, 1, 2, 4, 12 | SynCom9 community (2), exudates (3), root metabolites (1), shoot metabolites (1) | 30 mL | 3, 4, 5, S5, S6 – S15 |

**Table S7 Overview of library preparation and sequencing protocols applied for the different experiments in this study.**

**16S V5 – V7 gene amplicon PCR protocol**

| Experiment | Figures | DNA extraction kit | Sequencing chemistry | Median reads/sample | Rarefaction threshold |
|---|---|---|---|---|---|
| Microbiome in glass- vs clay-microcosms | 1, S1, S3 | QIAGEN® DNeasy® 96 PowerSoil® Pro Kit (Qiagen, Germany) | v2 500 cycles PE | 4228 | 2300 |

**Cycling conditions PCR1**

| Temperature | Time | cycles |
|---|---|---|
| 94 °C | 2 min | |
| 94 °C | 30 s | 25x |
| 55 °C | 30 s | |
| 72 °C | 30 s | |
| 72 °C | 5 min | |
| 15 °C | hold | |

**Primers used PCR1**

| Primer name | CS-linker | Frameshift | Primer sequence | Full sequence | Direction |
|---|---|---|---|---|---|
| CS1-799-F | ACACTGACGACATGGTTCTACA | | AACMGGATTAGATACCCKG | **ACACTGACGACATGGTTCTACAAACMGGATTAGATACCCKG** | foward |
| CS1_fs1_799F | ACACTGACGACATGGTTCTACA | T | AACMGGATTAGATACCCKG | **ACACTGACGACATGGTTCTACATAACMGGATTAGATACCCKG** | foward |
| CS1_fs2_799F | ACACTGACGACATGGTTCTACA | CT | AACMGGATTAGATACCCKG | **ACACTGACGACATGGTTCTACACTAACMGGATTAGATACCCKG** | foward |
| CS1_fs3_799F | ACACTGACGACATGGTTCTACA | ACT | AACMGGATTAGATACCCKG | **ACACTGACGACATGGTTCTACAACTAACMGGATTAGATACCCKG** | foward |
| CS1_fs4_799F | ACACTGACGACATGGTTCTACA | GACT | AACMGGATTAGATACCCKG | **ACACTGACGACATGGTTCTACAGACTAACMGGATTAGATACCCKG** | foward |
| CS2-1193-R | TACGGTAGCAGAGACTTGGTCT | | ACGTCATCCCCACCTTCC | **TACGGTAGCAGAGACTTGGTCTACGTCATCCCCACCTTCC** | reverse |
| CS2-fs1_1193-R | TACGGTAGCAGAGACTTGGTCT | T | ACGTCATCCCCACCTTCC | **TACGGTAGCAGAGACTTGGTCTTACGTCATCCCCACCTTCC** | reverse |
| CS2-fs2_1193-R | TACGGTAGCAGAGACTTGGTCT | CT | ACGTCATCCCCACCTTCC | **TACGGTAGCAGAGACTTGGTCTCTACGTCATCCCCACCTTCC** | reverse |
| CS2-fs3_1193-R | TACGGTAGCAGAGACTTGGTCT | ACT | ACGTCATCCCCACCTTCC | **TACGGTAGCAGAGACTTGGTCTACTACGTCATCCCCACCTTCC** | reverse |
| CS2-fs4_1193-R | TACGGTAGCAGAGACTTGGTCT | GACT | ACGTCATCCCCACCTTCC | **TACGGTAGCAGAGACTTGGTCTGACTACGTCATCCCCACCTTCC** | reverse |

**host-associated microbioma PCR (hamPCR)**

| Experiment | Figures | DNA extraction kit | Sequencing chemistry | Median reads/sample | Rarefaction threshold |
|---|---|---|---|---|---|
| Microbiome and exudate dynamics in glass-microcosms | 3, 4, 5, S5 B, S6 – S13 | QIAGEN® MagAttract® PowerSoil® DNA KF Kit (Qiagen, Germany) with KingFisher™ Flex™ Purification System (ThermoFisher Scientific, Waltham, MA) | v3 600 cycles PE | 20521 | 10500 |

**Cycling conditions PCR1**

| Temperature | Time | cycles |
|---|---|---|
| 94 °C | 2 min | |
| 94 °C | 30 s | 10x |
| 60 °C | 30 s | |

| | |
|---|---|
| 72 °C | 60 s |
| 72 °C | 10 min |
| 15 °C | hold |

**Primers used PCR1**

| Primer name | CS-linker | Frameshift | Primer sequence | Full sequence | Direction |
|---|---|---|---|---|---|
| CS1-799-F | ACACTGACGACATGGTTCTACA | | AACMGGATTAGATACCCKG | **ACACTGACGACATGGTTCTACAAACMGGATTAGATACCCKG** | foward |
| CS1_fs1_799F | ACACTGACGACATGGTTCTACA | T | AACMGGATTAGATACCCKG | **ACACTGACGACATGGTTCTACATAACMGGATTAGATACCCKG** | foward |
| CS1_fs2_799F | ACACTGACGACATGGTTCTACA | CT | AACMGGATTAGATACCCKG | **ACACTGACGACATGGTTCTACACTAACMGGATTAGATACCCKG** | foward |
| CS1_fs3_799F | ACACTGACGACATGGTTCTACA | ACT | AACMGGATTAGATACCCKG | **ACACTGACGACATGGTTCTACAACTAACMGGATTAGATACCCKG** | foward |
| CS1_fs4_799F | ACACTGACGACATGGTTCTACA | GACT | AACMGGATTAGATACCCKG | **ACACTGACGACATGGTTCTACAGACTAACMGGATTAGATACCCKG** | foward |
| CS1-At.GI-F | ACACTGACGACATGGTTCTACA | | CTGTAAAGATAAATGGGTCATCTAA | **ACACTGACGACATGGTTCTACACTGTAAAGATAAATGGGTCATCTAA** | forward |
| CS2-1193-R | TACGGTAGCAGAGACTTGGTCT | | ACGTCATCCCCACCTTCC | **TACGGTAGCAGAGACTTGGTCTACGTCATCCCCACCTTCC** | reverse |
| CS2-fs1_1193-R | TACGGTAGCAGAGACTTGGTCT | T | ACGTCATCCCCACCTTCC | **TACGGTAGCAGAGACTTGGTCTTACGTCATCCCCACCTTCC** | reverse |
| CS2-fs2_1193-R | TACGGTAGCAGAGACTTGGTCT | CT | ACGTCATCCCCACCTTCC | **TACGGTAGCAGAGACTTGGTCTCTACGTCATCCCCACCTTCC** | reverse |
| CS2-fs3_1193-R | TACGGTAGCAGAGACTTGGTCT | ACT | ACGTCATCCCCACCTTCC | **TACGGTAGCAGAGACTTGGTCTACTACGTCATCCCCACCTTCC** | reverse |
| CS2-fs4_1193-R | TACGGTAGCAGAGACTTGGTCT | GACT | ACGTCATCCCCACCTTCC | **TACGGTAGCAGAGACTTGGTCTGACTACGTCATCCCCACCTTCC** | reverse |
| CS2-At-GI-R | TACGGTAGCAGAGACTTGGTCT | | AAGGGTTCAGCTTTGTCAACAA | **TACGGTAGCAGAGACTTGGTCTAAGGGTTCAGCTTTGTCAACAA** | reverse |
